# Structural insights into Cas9-mediated prespacer selection in CRISPR-Cas adaptation

**DOI:** 10.1101/2025.06.12.659244

**Authors:** Ugne Gaizauskaite, Giedre Tamulaitiene, Arunas Silanskas, Giedrius Gasiunas, Virginijus Siksnys, Giedrius Sasnauskas

## Abstract

During CRISPR-Cas adaptation, prokaryotic cells become immunized by the insertion of foreign DNA fragments, termed spacers, into the host genome to serve as templates for RNA-guided immunity. Spacer acquisition relies on the Cas1-Cas2 integrase and accessory proteins like Cas4, which select DNA sequences flanked by the protospacer adjacent motif (PAM) and insert them into the CRISPR array. It has been shown that in type II-A systems selection of PAM-proximal prespacers is mediated by the effector nuclease Cas9, which forms a ‘supercomplex’ with the Cas1-Cas2 integrase and the Csn2 protein. However, the supercomplex structure and the role of the ring-like Csn2 protein remain unknown. Here, we present cryo-electron microscopy structures of the type II-A prespacer selection supercomplex in the DNA-scanning and two different PAM-bound configurations. Our study uncovers the mechanism of Cas9-mediated prespacer selection in type II-A CRISPR-Cas systems, and reveals the role of the accessory protein Csn2, which serves as a platform for the assembly of Cas9 and Cas1-Cas2 integrase on prespacer DNA, reminiscent of the sliding clamp in DNA replication. Repurposing of Cas9 by the CRISPR adaptation machinery for prespacer selection characterized here demonstrates Cas9 plasticity and expands our knowledge of the Cas9 biology.

## INTRODUCTION

In prokaryotes, CRISPR (clustered regularly interspaced short palindromic repeats) sequences and CRISPR-associated (Cas) proteins function as an adaptive immune system, protecting host cells from bacteriophages and other mobile genetic elements^1^. A CRISPR array consists of a leader sequence followed by conserved repeats interspersed with variable spacer sequences, originating from invading nucleic acids. During the adaptation stage of CRISPR-Cas immunity, new spacers are captured from the foreign DNA and inserted into the Leader-proximal end of the CRISPR array by the conserved Cas1-Cas2 integrase^2–6^. Upon insertion, spacers are used as templates for crRNA transcripts that serve as guides for CRISPR-Cas effector complexes, which recognize and destroy invading nucleic acids^7^.

In the DNA-targeting CRISPR-Cas systems, DNA destruction requires complementarity between crRNA and the target DNA, along with an adjacent PAM (protospacer adjacent motif) sequence, which is specifically recognized by the effector protein^8,9^. The PAM requirement in the interference stage dictates that each new spacer must be captured from an invading DNA fragment (prespacer) flanked by a PAM sequence, and inserted into the CRISPR array in a PAM-dependent orientation, but without the PAM sequence to prevent self-targeting^10^.

To date, various mechanisms for PAM-dependent prespacer selection and processing have been reported^11,12^. In many type I and some type V systems these functions are performed by the Cas4 endonuclease that is either associated or fused to the core integrase protein Cas1^13–19^. In type I-E systems the PAM-binding pocket is present in the Cas1 protein, enabling PAM-recognition and prespacer selection by the Cas1-Cas2 integrase complex^20–22^. Subsequent PAM removal and end-trimming is performed by accessory DnaQ-like endonucleases, Cas1 protein or host nucleases^20–25^. Notably, in the above systems, PAM-recognition during adaptation and interference steps is performed by separate proteins/complexes: the PAM sequence is recognized by Cas1/Cas4 during spacer acquisition, and by the Cascade/Cas12 effector complex during target DNA degradation. Conversely, in type II-A systems, PAM recognition during both adaptation and interference steps is carried out by the Cas9 effector nuclease^26,27^ that together with Cas1-Cas2 integrase and the auxiliary Csn2 protein^28^ are required for spacer acquisition^26,27^, and form a Cas9-Cas1-Cas2-Csn2 ‘supercomplex’^29–31^. Cryogenic electron microscopy (cryo-EM) analysis of complexes formed by the Cas1-Cas2 integrase and Csn2 from the *S. thermophilus* DGCC 7710 CRISPR3 (henceforth - St3) system has revealed several different assemblies consisting of Csn2 tetrameric rings bridged by Cas1-Cas2 integrases, however, the structure of the supercomplex remained to be established^30^.

Here we report cryo-EM structures of St3 Cas9-Cas1-Cas2-Csn2 supercomplex in the PAM search and two different PAM-bound configurations, which provide the structural basis for the functional coupling between Cas9 and spacer integration machinery, and reveal the multifaceted role of the accessory protein Csn2.

## RESULTS

### The St3 system

The type II-A St3 system from *S. thermophilus*^32^ consists of St3Cas9 effector nuclease, St3 Cas1-Cas2 integrase and the accessory protein St3Csn2 (Figure 1A). St3 CRISPR array comprises a leader sequence followed by 36 bp partially palindromic repeats interspersed by 30 bp spacer sequences. To uncover the molecular mechanism of spacer insertion in the St3 system (Figure 1B), we first used cryo-EM to determine structures of the Leader-Repeat1-specific St3 integrase and the PAM-specific St3Cas9 nuclease bound to the respective dsDNA fragments, followed by characterization of higher-order oligomeric complexes.

**Figure 1.**
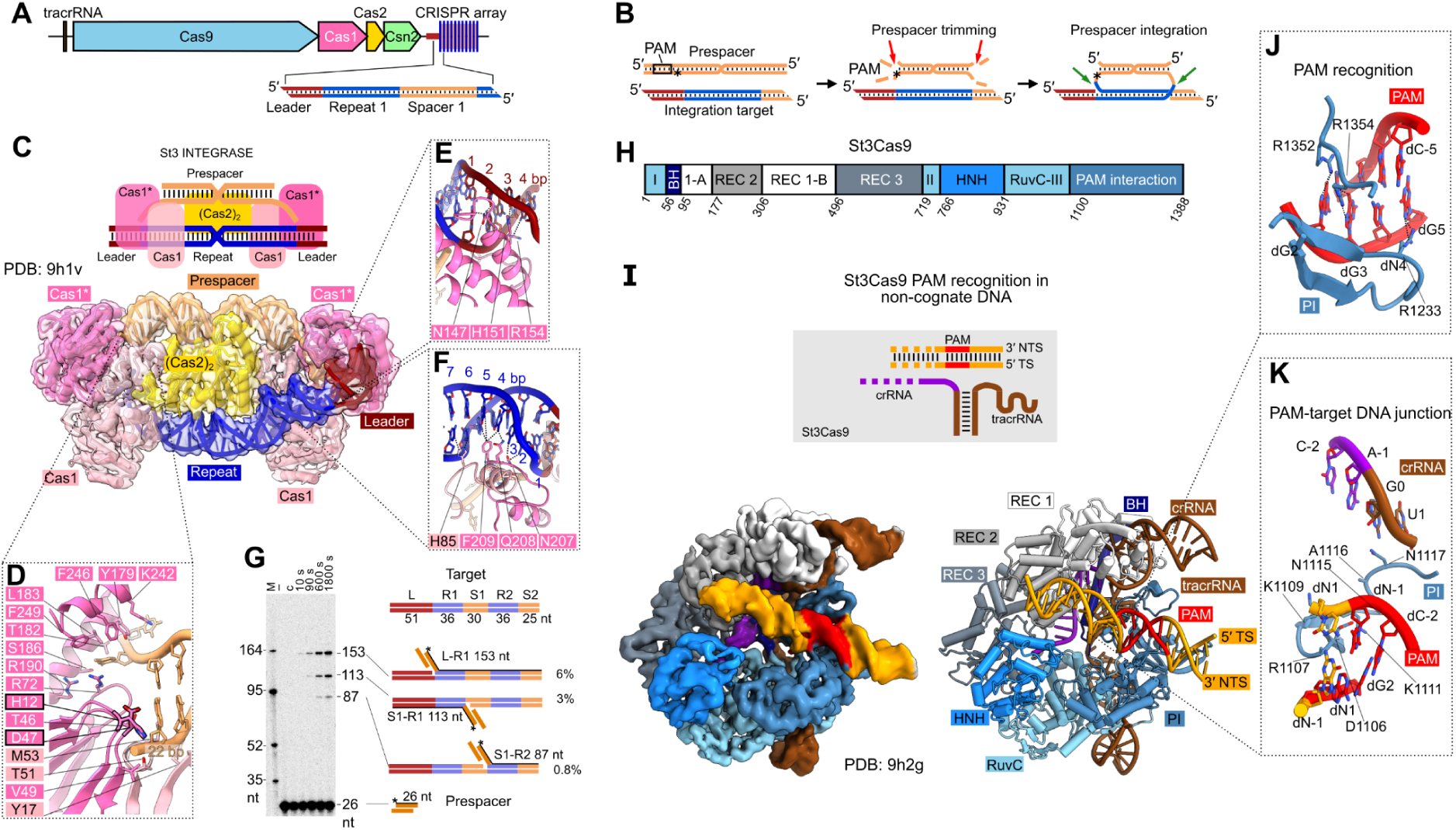
Sequence-specific components of the *S. thermophilus* CRISPR3 (St3) system. (A) Schematic representation of the *S. thermophilus* CRISPR3-Cas locus. (B) Prespacer acquisition in type II-A CRISPR-Cas systems. The asterisk (*) marks the PAM-proximal terminus of the prespacer. (C) Structure of the St3 Cas1-Cas2 integrase complex. Top: schematic representation of the (Cas1)_2_-(Cas2)_2_-(Cas1)_2_ complex with prespacer DNA (22 bp duplex region with 4 nt 3′-overhangs) and integration target (36 bp Repeat sequence surrounded by 7 bp Leader sequences); bottom: unsharpened semi-transparent cryo-EM map and atomic model. (D) Contacts of the integrase complex to the end of the 22 bp prespacer fragment. H12* and D47* residues stacking against the end of the 22bp prespacer fragment are encircled in black. (E-F) Recognition of the Leader-Repeat1 junction by the St3 integrase complex. Dotted lines mark H-bonds and van der Waals interactions. (G) St3 integrase preferentially integrates prespacers at the Leader-proximal Repeat sequence. Right: schematic representation of the integration target (CRISPR3 region with Leader and 2 repeat sequences), prespacer DNA (22 bp duplex with 4 nt 3′-overhangs) used in the reaction, and the reaction products detectable by denaturing-PAGE (left). Lanes 1-4 – integrase reactions with prespacer and integration target; lane ‘–’ prespacer only, lane ‘c’ – 1800 s reaction without the integration target, lane ‘M’ – DNA length markers. (H) St3Cas9 structural domains. (I) Structure of St3Cas9:crRNA:tracrRNA complex interacting with the PAM sequence in a DNA duplex lacking complementary to guide RNA. Top: schematic representation of the complex; center: unsharpened cryo-EM map; bottom: atomic model. (J-K) 5′-NGGNG-3′ PAM recognition and junction between PAM (red) and upstream (orange) DNA in the St3Cas9 ternary complex with the non-complementary DNA.

### Leader-Repeat recognition by the St3 Cas1-Cas2 integrase

Cryo-EM structures of St3 Cas1-Cas2 integrase bound to both prespacer and integration target DNA (Figure 1C) or to prespacer alone (Figure S1A) revealed similarities to the Cas1-Cas2 integrase from the *E. faecalis* (Ef) type II-A CRISPR-Cas system^5^ (Figure S1B). In the heterohexameric (Cas1)_2_-(Cas2)_2_-(Cas1)_2_ integrase, the inner non-catalytic Cas1 subunits interact with the central Cas2 homodimer, whereas the outer Cas1 subunits (henceforth - the catalytic subunits Cas1*) catalyze prespacer integration reactions at the opposite ends of the Repeat DNA (Figure 1B). The 30 bp prespacer is bound by the St3 integrase as an undistorted 22 bp DNA fragment with 4-nt 3′-overhangs at each end that are directed to the catalytic centers of the Cas1* subunits (Figure 1C). The 22 bp duplex length is defined by end-stacking of Cas1* H12* and D47* residues to both prespacer ends (Figure 1D). Prespacer DNA backbone forms numerous contacts to the positively charged residues of both Cas1/Cas1* and Cas2 proteins, a notable exception being divalent cation-mediated contacts to Cas2 D13 residues at the prespacer center (Figure S1C).

Concomitant binding to the Leader-Repeat DNA fragment has a minor effect on the overall conformation of the Cas1-Cas2 integrase (Figure S1D). As in the Ef Cas1-Cas2 complex^5^, St3 integrase inserts an α-helix (residues 148*-161*) of the catalytic Cas1* subunit into the minor groove of the Leader sequence, making contacts to its first 4 base pairs (Figure 1E). Direct readout of the 2-7 bp of the inverted repeat sequence of the CRISPR Repeat is mediated by loops from both the catalytic (residues 206*-211*) and the non-catalytic (residues 82-86) Cas1 subunits (Figure 1F). Despite promiscuous nature of the minor groove contacts to the bases in the Leader sequence, St3 integrase preferentially inserts new spacers at the Leader-end of a CRISPR array *in vitro*, with the most efficient reaction occurring at the Leader-Repeat1 junction (Figures 1G and S1E).

### PAM recognition by St3Cas9

St3Cas9 of the *S. thermophilus* CRISPR3 system, being among the first Cas9 proteins harnessed for genome editing applications^33^ and a close homolog of *S. pyogenes* (Sp) Cas9 (Figure S2A), requires a 5′-NGGNG-3′ PAM sequence immediately downstream of the target site^32^. Cryo-EM structure of the ternary complex of St3Cas9:crRNA:tracrRNA bound to a PAM and a cleaved target DNA (Figure S2B) revealed that St3Cas9 recognizes three guanines of the PAM sequence via three arginines of the PAM-interacting domain loop (residues 1220-1238, Figure S2A), whereas the strand-separation loop (residues 1105-1111) contacts the dsDNA terminus immediately upstream of PAM (Figure S2C).

During prespacer selection Cas9 has to recognize PAM in a DNA fragment lacking complementarity to the crRNA, therefore we have also determined the structure of St3Cas9:crRNA:tracrRNA bound to a DNA fragment carrying a PAM sequence but non-complementary to the crRNA. Only approx. half of St3Cas9:crRNA:tracrRNA complexes bound to such DNA retained the PAM-specific contacts (Figures 1H–1J) observed in the post-cleavage complex (Figure S2A), consistent with multi-micromolar affinity of Cas9 proteins to PAM-only DNA^34,35^. The remaining complexes formed loose non-specific interactions with the undistorted non-cognate DNA (Figure S2D), resembling the protein-DNA interaction observed in SpCas9 complex with the unbent non-complementary DNA (PDB: 7s3h^34^). In the PAM-recognizing complexes, St3Cas9 inserts the strand-separation loop between the PAM and the upstream DNA, thereby unstacking the respective base pairs and distorting the DNA (Figure 1K). Overall conformation of this St3Cas9:crRNA:tracrRNA:DNA assembly matches that of the DNA-free St3Cas9:crRNA:tracrRNA complex (Figures S2E–S2G).

### Structures of St3 supercomplex bound to DNA

To uncover how St3Cas9 participates in the adaptation process, we have reconstituted the St3 supercomplex by mixing purified components of the St3 system (the pre-assembled St3Cas9:crRNA:tracrRNA complex, St3Csn2 and St3 Cas1-Cas2 integrase) with a 49 bp PAM-containing dsDNA fragment with no complementarity to crRNA. The supercomplex stoichiometry is consistent with 1:1:1:1 assembly of Cas9-RNA, Cas1-Cas2 heterohexamer, Csn2 tetramer, and dsDNA (Figures S3A–S3D); no supercomplex was formed in the absence of dsDNA (Figures S3E–S3F). Although the supercomplex constituted the minor fraction of the cryo-EM sample (<25%, see Methods), presumably due to low stability, we were able to determine its structures with three different DNA conformations to 3.39-3.72 Å resolution (Figures 2A–2D). Cryo-EM maps allowed us to resolve one Cas9-RNA complex, a single St3Csn2 tetramer, part of a heterohexameric Cas1-Cas2 integrase (Cas1 dimer with a bound Cas2 fragment), and approx. 40 bp dsDNA. In all three structures one DNA end is encircled by the homotetrameric Csn2 ring (Figures 2B–2D and Movie S1). The opposite DNA end in the PAM-unbound structure (supercomplex in search conformation) loosely associates with St3Cas9 (Figure 2B). In the two PAM-bound structures, the PAM forms sequence-specific contacts with the St3Cas9 PAM-interacting domain, whereas the strand separation loop intercalates between the PAM and the upstream DNA, similarly to the structure of isolated St3Cas9 with PAM-only DNA (Figures 2C–2D and 1K). In the first PAM-bound structure (the docked conformation), the PAM-upstream DNA is directed towards the Csn2 ring by circumventing the Cas9 REC2 domain, resulting in DNA bending (Figure 2C). In the second PAM-bound structure (the locked conformation), the REC2 domain is repositioned (Figure 2E and Movie S2), opening a straight path for the DNA from the PAM recognition domain to the Csn2 ring (Figure 2D). Identical protein conformations, and overlapping DNA paths in the search and docked structures (Figures 2B–2D) suggest that the docked complex is an intermediate formed during transition of the PAM-unbound (search) supercomplex to the PAM-bound locked conformation with a displaced REC2 domain and well resolved straight DNA (Movie S2).

**Figure 2.**
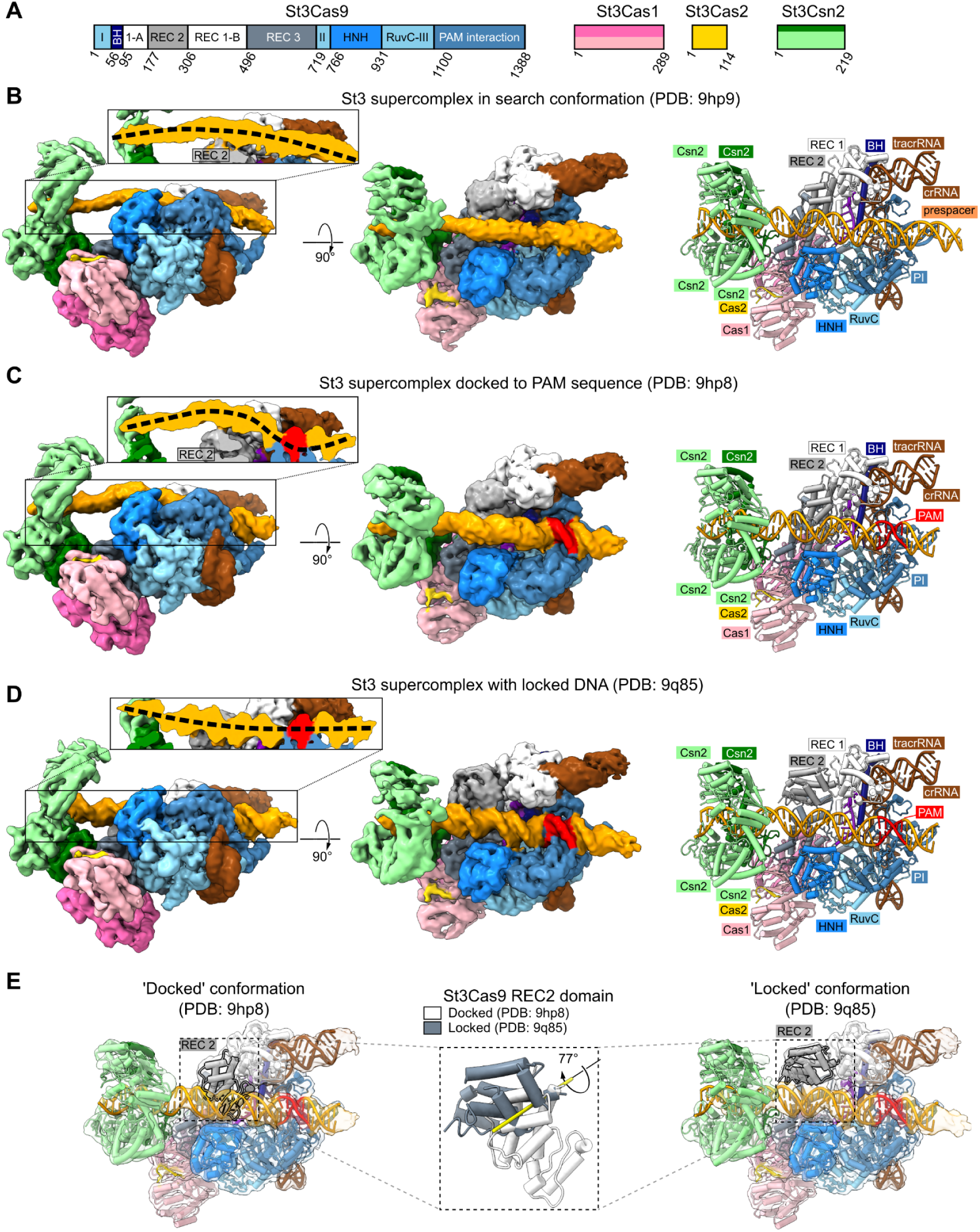
The supercomplex of the *S. thermophilus* CRISPR3 (St3) system. (A) Domain organization and coloring scheme of the St3 system components. (B) St3 supercomplex forming non-specific interactions with the DNA. (C) St3 supercomplex docked to PAM sequence. (D) PAM-bound St3 supercomplex in the locked conformation. (E) Movement of St3Cas9 REC2 domain during transition of the PAM-bound ‘docked’ supercomplex into the ‘locked’ conformation. In panels (B)-(D), different views of the unsharpened cryo-EM maps are shown, the insets depict slices of the respective maps along the captured DNA in the regions marked by black rectangles; the overall paths of the DNAs bound to the supercomplexes are indicated by dashed lines. Atomic models of the respective complexes are shown on the right. Coloring in panels (B)-(E) follows the scheme in panel (A). Cas1 and Cas1* denote the non-catalytic and catalytic Cas1 subunits, respectively. The Csn2 subunit that forms interactions with Cas1/Cas1* and Cas9 proteins is colored in green, the remaining three Csn2 subunits are colored in pale green.

### Protein interfaces within the supercomplex

In the St3 supercomplex we observe 3 interfaces between the Cas9, Csn2 and integrase proteins (Figures 3A–3D and S4A): (1) both subunits of the Cas1 homodimer interact with a single subunit of the Csn2 tetramer; this 619 Å^2^ interface is formed by the N- and C-termini of Csn2, and the N-terminal regions of Cas1 and Cas1*; (2) both subunits of the Cas1 homodimer interact with the Cas9 REC3 domain (637 Å^2^); (3) REC3 domain of Cas9 interacts with the N-terminal region of the same Csn2 subunit as Cas1 (351 Å^2^). Interface 1 (Figure 3B) was previously observed in the St3 Csn2-integrase ‘monomer’ assembly (PDB: 6qxt, Figures S4B–S4C)^30^, in which St3 integrase adopts a similar conformation as in the supercomplex (Figure S4D), with its prespacer DNA binding groove perpendicular to the captured DNA fragment (Figure S4E). Lack of the second Csn2 ring in the supercomplex likely renders the second Cas1 dimer mobile and therefore unresolved in our structure (Figure S4B). Formation of Interfaces 1-2 occludes some of the integrase residues that in the integration-ready conformation contact the prespacer and Cas2 (Figure 3E), thus locking it in an inactive state. The smallest Interface 3 brings together two Cas9 REC3 domain α-helices and the N-terminal Csn2 residues, some of which also contact Cas1 (Figures 3D and S4A). To test the importance of Interface 1 and Interface 2 in the spacer acquisition in the native host without interfering with the Cas1-Cas2 integrase function, we have constructed the ΔCRISPR1ΔCRISPR4 variant of the *S. thermophilus* DGCC 7710 strain retaining only the St3 CRISPR-Cas system, and introduced into its genome *St3csn2* Y212R+I213R+M215R mutations targeting the Cas1-Csn2 interface (Interface 1) or *St3cas9* Q544A+K706E mutations targeting the Cas9-Cas1 interface (Interface 2), but not compromising the mutant protein stability nor St3Cas9 DNA cleavage activity *in vitro* (Figures S5A–S5B). We have found that Interface 1 mutations blocked formation of the supercomplex *in vitro* (Figure S3G), and both sets of mutations either completely (Interface 1) or almost completely (Interface 2) abolished acquisition of new spacers when the respective *S. thermophilus* strains were challenged with the D2972 phage (Figure 3F), confirming the relevance of the observed supercomplex structure in spacer acquisition.

**Figure 3.**
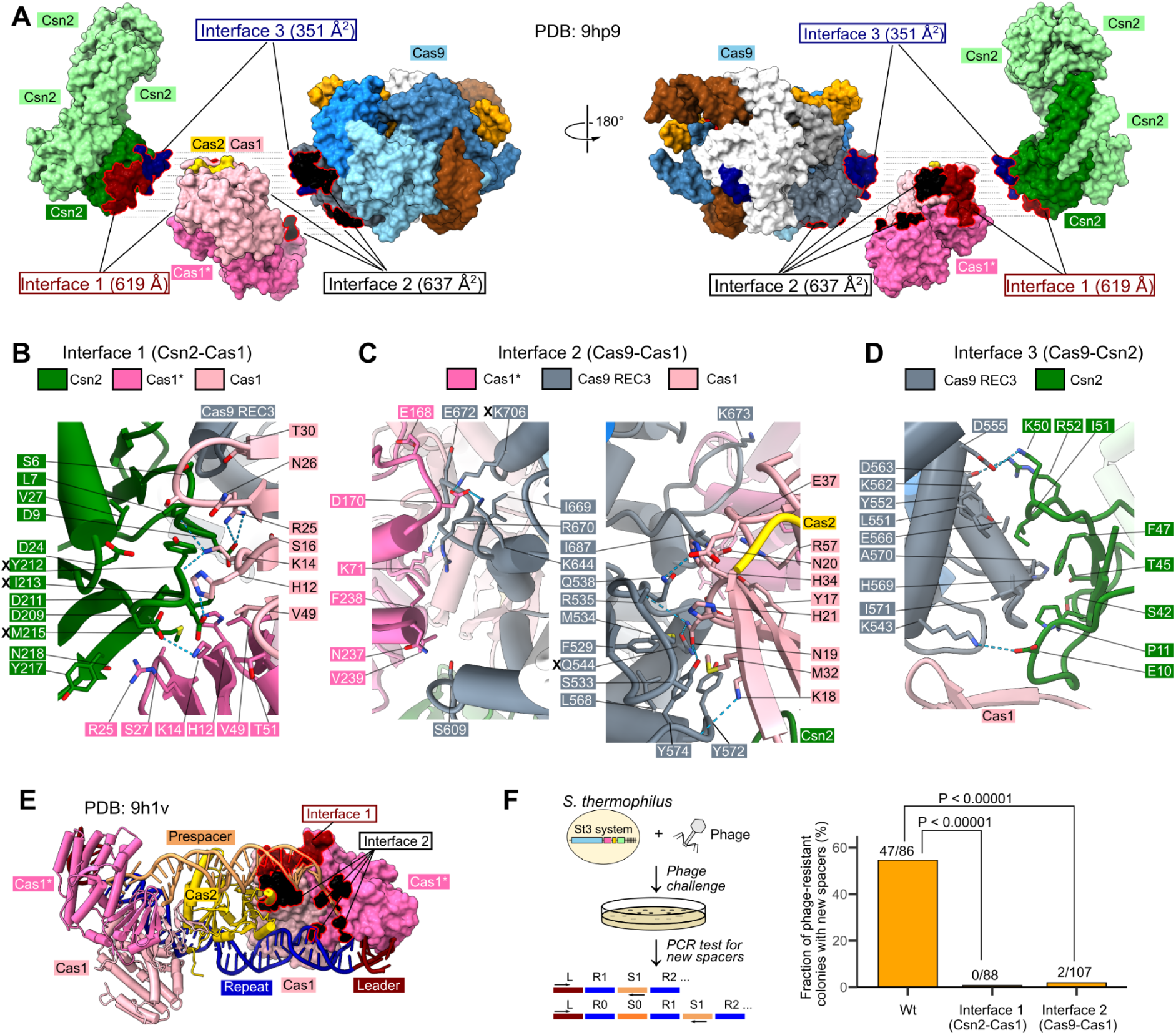
Protein-protein interfaces in the St3 supercomplex. (A) Different views of Interface 1 (Csn2-(Cas1)_2_, maroon), Interface 2 (Cas9-(Cas1)_2_, black) and Interface 3 (Csn2-Cas9, navy). (B)-(D) Contacts between Csn2, Cas1* (catalytic Cas1 subunit), Cas1 (non-catalytic Cas1 subunit) and Cas9 REC3 domain in the St3 supercomplex. Interface 1 and 2 residues mutated in this study are marked by ‘X’. Zoom-in views are provided in Figure S4A. (E) Cas1 residues that form Interface 1 (maroon) and Interface 2 (black) in the St3 supercomplex, during the integration reaction interact with the prespacer, the adjacent Cas2 subunit, and the integration target. (F) Effect of Interface 1 and Interface 2 mutations (St3Csn2 triple mutant St3Csn2^Y212R+I213R+M215R^ and St3Cas9 double mutant St3Cas9^Q544A+K706E^, respectively) on spacer acquisition by the St3 system *in vivo*. Left: *S. thermophilus* strain carrying St3 system (WT or Interface 1/2 mutant) was challenged with the D2972 phage, and the isolated bacteriophage insensitive clones were tested for spacer acquisition. Right: the bar graph showing the fraction of clones that acquired new spacers. Total number of tested colonies is indicated above the bars. The p values were calculated using a two proportion Z-test as described in the Methods section.

### Supercomplex as an integrase recruitment platform

The supercomplex characterized here is not compatible with the prespacer integration reaction, as both termini of the captured prespacer DNA are inaccessible, the prespacer-binding groove of the Cas1-Cas2 integrase in the supercomplex is perpendicular to the captured prespacer (Figure S4E), and some Cas1 DNA binding surfaces are occluded (Figure 3E). This conclusion is further supported by the inhibition of the St3 integration reactions by St3Csn2 (Figure S5C).

Noteworthy, reactions of an isolated St3 integrase *in vitro* are limited to integration of pre-processed prespacers of correct length in a PAM-independent manner (Figure S5C). In contrast, prespacer acquisition *in vivo* depends on both St3Cas9 and St3Csn2^1^, and ensures selection, processing and insertion of prespacers in a PAM-defined orientation. We therefore propose that the supercomplex structures described above represent the first steps in spacer acquisition in host cells, i. e. capturing a DNA fragment and selecting its PAM-containing region prior to the integration reaction. The PAM-unbound complex presumably represents the search process, where the supercomplex, upon assembling on an open-end dsDNA fragment, slides along the DNA probing for a valid PAM sequence. Recognition of the correct PAM by Cas9 docks it onto the PAM-containing DNA fragment (the docked complex), followed by its transition to the locked conformation (Movie S2). Notably, most of the PAM-proximal 30 bp would-be-spacer DNA captured in the locked supercomplex is exposed (Figure 4A), providing a probable recruitment platform for a separate integrase complex responsible for catalysis, or capture of the mobile part of the integrase already present in the supercomplex that was not resolved in the structures (a full Cas1 dimer and most of a Cas2 dimer).

**Figure 4.**
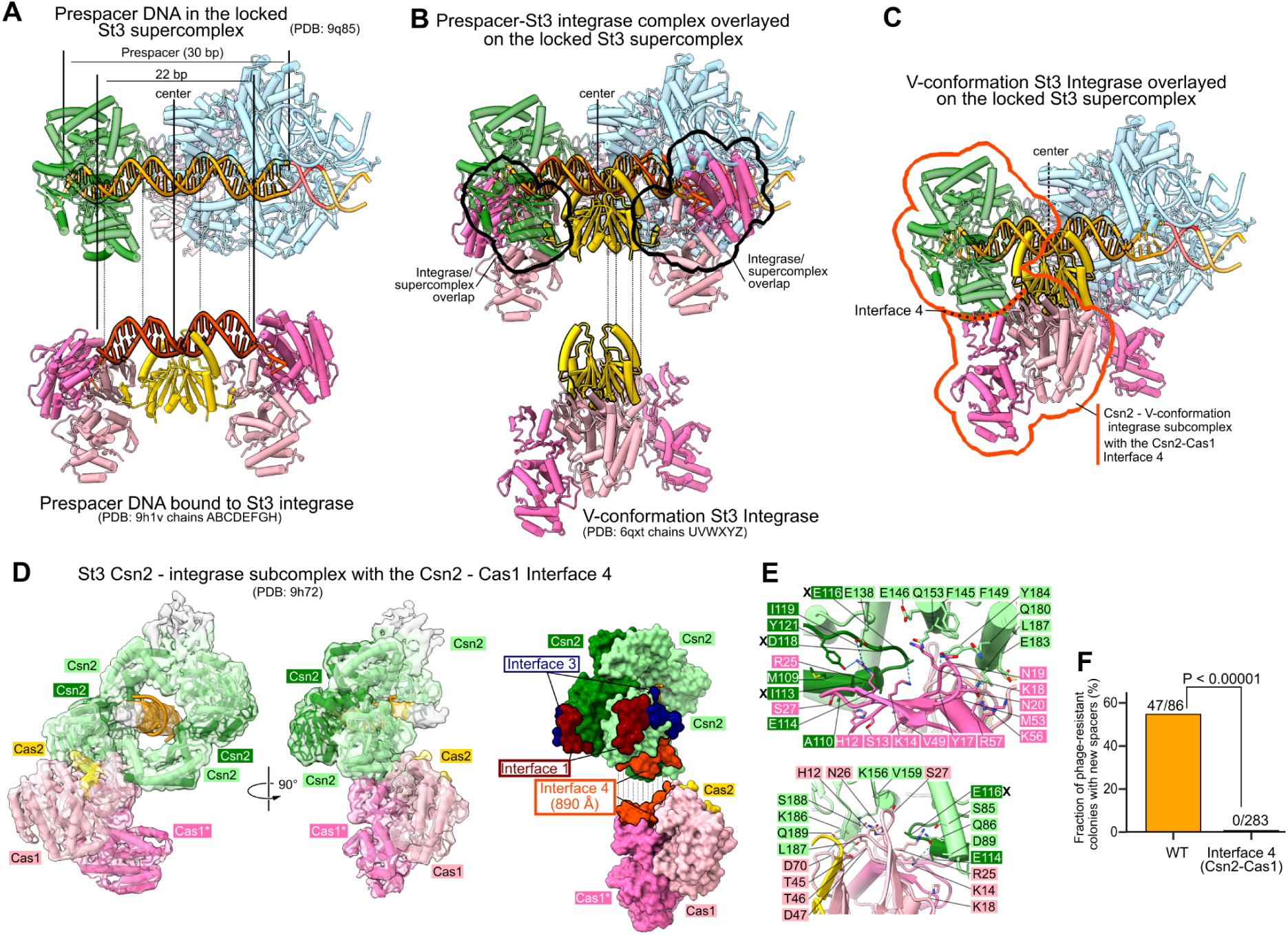
Supercomplex as a Cas1-Cas2 integrase recruitment platform. (A) Prespacer DNA captured in the ‘locked’ St3 supercomplex and bound to Cas1-Cas2 integrase. Boundaries of the 30 bp would-be-spacer DNA, its center, and the 22 bp central part recognized by the integrase as dsDNA are indicated. (B) An overlay of the prespacer bound to St3 integrase on the prespacer in the supercomplex places the integrase on the exposed face of the DNA. Black outlines indicate steric clashes between Cas1 subunits of the overlaid integrase and Csn2/Cas9 in the supercomplex. The St3 integrase V-conformation is shown below the overlay. Dotted lines in panels a-b connect the overlaid structural elements. (C) An overlay of V-conformation St3 integrase on the prespacer DNA in the supercomplex, places one Cas1 dimer in the vicinity of Csn2, forming a Csn2 - integrase subcomplex (orange-red outline) with the alternative Cas1-Csn2 interface (Interface 4). (D) Left: cryo-EM structure of the Csn2 - integrase subcomplex forming the Interface 4. Right: St3Csn2 and St3Cas1 Interface 4-forming residues (orange). Interface 1 (maroon) and Interface 3 (navy) residues are also shown on both faces of the Csn2 tetramer. (E) Contacts between two adjacent Csn2 subunits and Cas1 homodimer at the Interface 4. Contacts to the catalytic and the non-catalytic Cas1 subunits are shown in the upper and lower subpanels, respectively. Coloring of Csn2 and integrase subunits follows the coloring scheme in (D). Residues mutated in this study are marked by ‘X’. (F) Effect of Interface 4 mutations (St3Csn2 triple mutant St3Csn2^I113R+E116R+D118R^) on spacer acquisition by the St3 system in the native *S. thermophilus* cells. Phage challenge experiments were performed as in Figure 3F. Data for the WT St3 system is also shown in Figure 3F. The p values were calculated using a two proportion Z-test as described in the Methods section.

Indeed, an overlay of the prespacer-bound St3 integrase on the DNA captured in the locked supercomplex suggests that the recruited integrase must engage prespacer DNA in the supercomplex from the exposed side (Figures 4A–4B). A similar structural context for integrase/Csn2 ring/dsDNA interaction is observed in the previously reported St3 integrase-Csn2 ‘dimer’ assembly (PDB: 6qxt, 8.9 Å resolution)^30^, where two compact V-conformation integrase complexes contact dsDNA via the Cas2 subunits, whereas Cas1 dimers interact with the adjacent (Csn2)_4_ rings (Figure S6A). Remarkably, positioning a V-shaped integrase so that the Cas2 dimer faces the center of the 30 bp locked would-be-spacer DNA 15 bp upstream of PAM, places the Csn2-proximal Cas1 dimer adjacent to the Csn2 ring of the supercomplex, forming an alternative Cas1-Csn2 interface observed in the ‘dimer’ complex (Figures 4C and S6A, henceforth referred to as Interface 4). Notably, the relative positioning and orientation of the integrase region resolved in the supercomplex structure (comprising a Cas1 dimer with a Cas2 fragment), and the modeled Cas1-Cas2 subunits of the V-shaped integrase docked on the supercomplex, are also compatible with a mechanism in which Interface 4 and interactions with the exposed prespacer are formed by the mobile (Cas1)_2_-(Cas2)_2_ part of the integrase already present in the supercomplex (Figure S6B).

The likely transient nature of this hypothesized supercomplex, involving the newly recruited or rearranged integrase complex, precluded its structural characterization. However, we were able to study the Interface 4 (total area approx. 890 Å^2^) in greater detail by determining a 3.4 Å cryo-EM structure of St3Csn2^Y212R+I213R+M215R^ (St3Csn2 variant with disabled Interface 1) in complex with St3 integrase (a single Cas1 dimer and fragment of Cas2 resolved) and a dsDNA molecule (Figures 4D and S6C). Interface 4 is formed by both Cas1 subunits, ‘head’ and ‘tail’ residues of one Csn2 subunit, and the ‘tail’ residues of the adjacent Csn2 subunit (Figures 4D–4E and S7A). Csn2 Interface 4 residues do not overlap with Csn2 Interface 1 (Csn2-Cas1) and Interface 3 (Csn2-Cas9) residues (Figures 4D and S7A), enabling direct assessment of its importance. Introducing Interface 4-disrupting St3*csn2* mutations I113R+E116R+D118R, which did not compromise St3Csn2 stability (Figure S5B) and did not eliminate supercomplex formation *in vitro* (Figure S3H), into the StΔCRISPR1ΔCRISPR4 genome abolished acquisition of new spacers (Figure 4F). Thus, our data shows that St3Csn2 can interact with the integrase in two different ways - via Interface 1 or Interface 4, both of which are critical for spacer acquisition (Figures 3F and 4F), - lending support to the proposed integrase recruitment/rearrangement mediated by Interface 4, as depicted in Figure 4C and S6B.

## DISCUSSION

Our findings provide the structural mechanism for prespacer selection employed by the type II-A CRISPR-Cas systems. Unlike the previously characterized type I systems, where spacer selection, trimming and integration steps occur in the context of the Cas1-Cas2 integrase^13,15,20,21^, the St3 type II-A system characterized here employs Cas9-dependent prespacer selection within the supercomplex, which allows the CRISPR system to re-use Cas9 for both prespacer capture during adaptation, and target recognition during interference. This also enables selection of prespacers with long PAM sequences, characteristic to some type II-A Cas9 proteins, as opposed to PAM readout by Cas1/Cas4 in the type I systems, which is limited to a few nucleotides in ssDNA^13,15,20,21^.

Prespacer selection within the supercomplex must be followed by its integration into the CRISPR array. Noteworthy, type I CRISPR-Cas systems first integrate the PAM-distal end of the selected prespacer at the Leader-Repeat1 junction, the preferred integration site, due to occlusion of the PAM-proximal end by the PAM-bound Cas1 or Cas4 subunits (Figure S8)^13,15,20,21^. However, spacer orientation is reversed for the Cas9-encoding type II-A systems^11^, which insert the PAM-proximal prespacer end at the optimal Leader-Repeat1 junction, and the PAM-distal end at the Repeat1-Spacer1 junction (Figures 1B and S8). In addition to retention of the correct prespacer directionality respective to PAM, the spacer acquisition process in the type II-A St3 system must involve the Cas9/Cas1/Csn2 interfaces characterized in the present study, and ensure PAM removal/prespacer trimming to the correct length of 30 bp.

A hypothetical spacer acquisition mechanism that fulfills the above criteria, but requires further experimental validation, is depicted in Figure 5. Supercomplex assembly on open-end DNA in the search conformation (#1), its docking to a PAM sequence and transition to the locked conformation (#2–#3) form an integrase recruitment platform. Integrase binding to prespacer DNA captured in the locked supercomplex via Interface 4 (#4, reminiscent of the overlays shown in Figures 4C and S6B) may initiate the prespacer hand-off. A stepwise mechanism, where in the first step the Cas9-proximal integrase half displaces Cas9 and captures the PAM-proximal prespacer end (#5), making the end available for host nucleases to generate an integration-ready 4-nt 3′ overhang at the PAM-proximal prespacer end, would direct its integration to the Leader-Repeat1 junction of the captured CRISPR region (#6), the preferred integration site (Figure 1G). Integration of one prespacer end by an asymmetric integrase is consistent with the presence of asymmetric St3 integrase complexes bound to a half-integration product formed *in situ* in integrase samples containing prespacer and target DNA (Figures S1F–S1H). Completion of the reaction would then require release of the PAM-distal prespacer terminus by Csn2, followed by its capture by the PAM-distal part of the integrase, end processing and integration at the remaining Repeat1-Spacer1 junction (#7).

**Figure 5.**
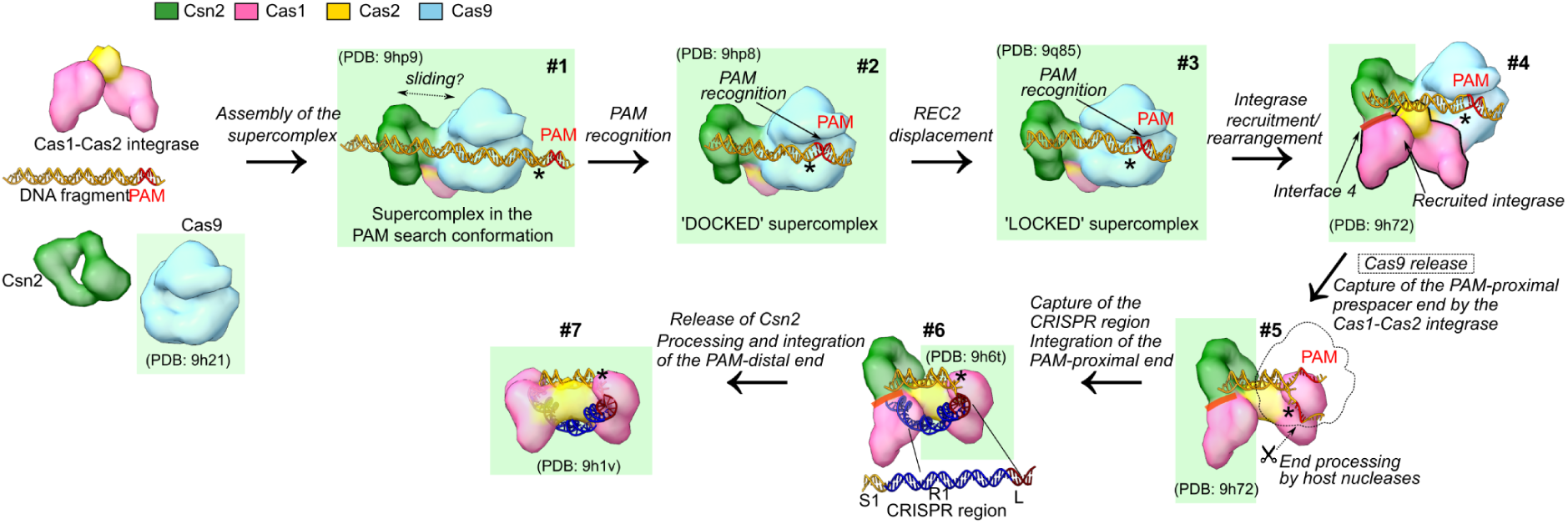
Hypothetical spacer acquisition mechanism in a type II-A St3 system. Complexes and subcomplexes structurally characterized in this study are marked by a green background.

Though the exact spacer acquisition mechanism for type II-A systems remains to be determined, a pivotal role in this process is carried out by the type II-A-specific accessory protein Csn2. First, Csn2 provides a platform for supercomplex formation, which selects a prespacer flanked by a PAM sequence, and proposed recruitment/rearrangement of the Cas1-Cas2 integrase, which takes over the captured prespacer from the supercomplex. Second, Csn2 ring protects the PAM-distal prespacer end, thereby directing its integration to the Repeat1-Spacer1 site (Figure 5). The ability of Csn2 to encircle dsDNA and slide on it^36^ makes its role in spacer acquisition reminiscent of the sliding clamp, which serves as a versatile platform for the recruitment and coordinated action of numerous factors involved in DNA replication, repair, and chromatin assembly^37^.

The architecture of the St3 spacer acquisition complexes is likely preserved in other type II-A systems, such as Sp and Ef, since structural elements involved in the intersubunit contacts between Cas9/Csn2/Cas1 are conserved, despite limited sequence similarity (Figure S7). This conservation is further supported by the deep-scanning mutagenesis of Sp CRISPR-Cas proteins^38^ and the reported MW of Sp supercomplex, which is consistent with 1:1:1 Cas9-RNA:integrase:(Csn2)_4_ assembly^31^. Spacer acquisition in the Cas9-encoding type II-B and II-C systems must be different due to the absence of Csn2 (II-C) or the presence of Cas4 (II-B). Although prespacer selection mechanism for these systems also remains to be determined, the overall similarity of type II-B integration machinery (including the presence of Cas4) to type I systems^39^, and short ‘NGG’ PAM sequences recognized by II-B Cas9 nucleases^40–42^, suggests selection of PAM-proximal prespacers by Cas4 in the context of Cas1-Cas2 integrase, whereas more diverse and often longer PAM sequences recognized by the type II-C Cas9 proteins^40^ imply that Cas9 itself functions as the PAM recognition module. Moreover, the absence of Csn2 in type II-C systems, and consequently the lack of PAM-distal end protection, may explain its integration into the Leader-Repeat1 site in a reverse orientation to type II-A systems (Figure S8), which is compensated by the opposite transcription direction of type II-C CRISPR arrays relative to the leader sequence^43,44^.

CRISPR effector proteins and complexes have been repurposed by nature for different functions, e.g., for regulation of CRISPR-Cas expression^45^, or as specificity modules in bacterial CRISPR-associated transposons^46–48^. Our findings add a new dimension to this general theme showing how Cas9 endonuclease, which provides DNA interference^49,50^, was harnessed by the CRISPR adaptation machinery for prespacer selection. We anticipate that the structures presented here will pave the way for further development of technologies that rely on the CRISPR spacer acquisition for molecular recording and information storage^51^.

## METHODS

### *S. thermophilus* genome editing

To secure CRISPR3 (St3) as the only active CRISPR-Cas system of the *S. thermophilus* DGCC 7710 strain, CRISPR1 (type II-A) and CRISPR4 (type I-E) systems were eliminated from the bacterial genome via homologous recombination^52^. ΔCRISPR1 and ΔCRISPR4 deletion cassettes were constructed by joining together 1 kbp upstream and 1 kbp downstream homology arms of the respective systems using overlap extension PCR (all oligonucleotide sequences used in this study are listed in Table S1), which was performed with Phusion DNA polymerase (Thermo Fisher Scientific). Each deletion cassette was transformed into naturally competent *S. thermophilus* DGCC 7710 cells as described previously ^53^. For the transformation, 3.3 µg of linear DNA and 1 µM of the competence-stimulating peptide ComS_17–24_ (sequence LPYFAGCL, synthesized by Peptide 2.0, purity of >97%) were added to 100 µL of *S. thermophilus* cells, which were grown in the chemically-defined medium (CDM). Transformed cells were 1000× diluted and plated on LM17 agar plates. Resulting colonies were restreaked on fresh LM17 agar plates in order to remove contaminating DNA. Colonies with successfully deleted CRISPR-Cas systems were identified by colony PCR using DreamTaq DNA polymerase (Thermo Fisher Scientific). CRISPR-Cas knockouts were performed consecutively resulting in the double-deletion mutant StΔCRISPR1ΔCRISPR4, which was further used for mutagenesis by introducing point mutations into the St3 system.

St3 point mutations were introduced into the StΔCRISPR1ΔCRISPR4 genome using the same method as mentioned before, the only difference being that instead of deletion, genomic DNA regions were substituted. Bacterial cells were transformed by a linear DNA fragment containing the point mutations of interest as well as silent mutations creating new restriction sites, which were necessary for colony screening. For Interface 1 (St3Csn2 Y212R+I213R+M215R) and Interface 4 (St3Csn2 I113R+E116R+D118R) mutagenesis, the linear DNA fragment was obtained by amplifying and assembling together 3 pieces of DNA using overlap extension PCR (one overlap introduces Csn2 mutations of interest, another overlap adds the restriction site for Eco88I). The final 2 kbp long PCR product was subcloned into pJET1.2 vector for Sanger sequencing and then again amplified by PCR to be used for transformation. For Interface 2 (St3Cas9 Q544A+K706E) mutagenesis, site directed mutagenesis of a whole pBAD plasmid encoding St3Cas9_C-His gene was performed via PCR in two rounds to consecutively introduce both point mutations and their respective silent mutations (an additional Bsp119I restriction site in the case of Q544A, a disrupted DraI restriction site in the case of K706E). The mutations were confirmed by Sanger sequencing, and the 2 kbp long *cas9* gene region encompassing all mutations was PCR-amplified for transformation.

PCR fragments containing interface mutations of interest were transformed into the naturally competent *S. thermophilus* cells as described above. After the transformation, colonies carrying the desired mutations of St3 protein interfaces 1, 2 and 4 were selected by restriction of colony PCR products. Subsequently, the DNA region covering the homologous recombination fragment and additional 1 kbp upstream and downstream DNA fragments of the selected *S. thermophilus* colonies were amplified by PCR and sequenced by the Sanger method in order to confirm the mutations of interest and to ensure the integrity of the St3 system.

### Isolation and analysis of bacteriophage insensitive mutants (BIMs)

*S. thermophilus* mutant strains were subjected to the phage challenge assay as described previously ^54^. Briefly, each strain was grown in LM17 medium at 42 °C until the mid-log phase (optical density at 600 nm (OD_600_) 0.6 to 0.8). Then, lytic D2972 bacteriophage and 100 µL of the cell culture were added to 3 mL of molten LM17 soft agar supplemented with 10 mM CaCl_2_ to achieve a multiplicity of infection (MOI) of 0.5. Soft agar was poured on top of LM17 agar plates with 10 mM CaCl_2_, and incubated at 42 °C for 24-36 h. Surviving colonies (CRISPR and non-CRISPR bacteriophage insensitive mutants, BIMs) were streaked on LM17 agar plates and screened for spacer acquisition in the St3 CRISPR locus by PCR using primers CR3fwd and CR3rev. StΔCRISPR1ΔCRISPR4 strain was used as a WT control. With each *S. thermophilus* strain, the phage challenge assay was performed repeatedly until the sample size exceeded approximately 80 BIMs. The proportions of BIMs containing new prespacers for the WT and mutant St3 systems were compared using the Z-test. Z values were calculated using the formula (1):

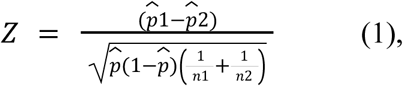

where *p̂*1 and *p̂*2 are the two sample proportions being compared, n1 and n2 are the two sample sizes, and *p̂* is the overall sample proportion.

### Protein expression

St3Cas9 (C-terminal HisTag, vector pBAD, described in^50^) and its mutant variant Cas9^Q544A+K706E^ were overexpressed in *E. coli* DH10B strain with L-arabinose (0.2% w/v) as an inductor. Expression of St3Csn2 (N-terminal HisTag, vector pACYCDuet-1, this study) and its mutant variant St3Csn2^Y212R+I213R+M215R^ was induced with IPTG (final concentration 0.2 mM) in the *E. coli* BL21 (DE3) strain. To purify St3 Cas1-Cas2 integrase, Cas1 (C-terminal HisTag, vector pETDuet-1, construct described in^30^, or C-terminal StrepII-Tag, vector pETDuet-1, this study) was co-expressed with untagged Cas2 (pACYC184-based plasmid St3Δ*cas9*Δ*cas1*, described in^55^) upon induction with 0.2 mM IPTG in the *E. coli* BL21 (DE3) strain. In each case, recombinant protein expression was performed as follows: cells harboring the plasmids of interest were grown at 37 °C in LB medium supplemented with appropriate antibiotics (100 μg/ml of ampicillin and/or 30 μg/ml of chloramphenicol) until the optical density at 600 nm (OD_600_) reached 0.5. Then, protein expression was induced by adding the appropriate inductor, followed by incubation at 16 °C for 16-18 h.

### Protein purification

After protein expression, all cells containing proteins of interest were collected by centrifugation, resuspended in Buffer 1 (20 mM Tris-HCl (pH 8.0 at 25 °C), 1 M NaCl, 5 mM 2-mercaptoethanol, 2 mM phenylmethylsulfonyl fluoride) and disrupted by sonication. After removing cell debris by centrifugation, the supernatants with WT St3Cas9 (C-terminal HisTag), St3Cas9^Q544A+K706E^ (C-terminal HisTag) and St3 Cas1-Cas2 integrase (C-terminal HisTag on St3Cas1) proteins were loaded on a Ni^2+^-charged HiTrap chelating HP column (Cytiva) and proteins were eluted with a linear gradient of increasing imidazole concentration (from 25 to 500 mM) in Buffer 2 (20 mM Tris-HCl (pH 8.0 at 25 °C), 500 mM NaCl, 5 mM 2-mercaptoethanol). The supernatant with St3 integrase complex (C-terminal StrepII-Tag on St3Cas1) was loaded onto a StrepTrap column (Cytiva), washed with Buffer 2, and eluted with 2.5 mM D-desthiobiotin solution. Fractions containing the target proteins were pooled, diluted with Buffer 3 (20 mM Tris-HCl (pH 8.0 at 25 °C), 5 mM 2-mercaptoethanol, 10% (v/v) glycerol) to decrease NaCl concentration until 250 mM, loaded on a HiTrap Heparin HP column (Cytiva) and eluted with a liner gradient of increasing NaCl concentration from 250 to 1000 mM.

The supernatants containing WT St3Csn2 protein (N-terminal HisTag) and its mutant variant St3Csn2^Y212R+I213R+M215R^ were loaded on a Ni^2+^-charged HiTrap chelating HP column (Cytiva) and eluted with a linear gradient of increasing imidazole concentration (from 25 to 500 mM) in Buffer 2. Fractions containing proteins of interest were collected, concentrated and loaded on the HiLoad 16/600 Superdex 200 gel-filtration column (Cytiva) equilibrated with Buffer 2.

Fractions with purified proteins were pooled and dialysed against a buffer containing 20 mM Tris-HCl (pH 8.0 at 25 °C), 500 mM NaCl, 2 mM dithiothreitol (DTT) and 50% (v/v) glycerol.

### Prespacer integration assays

Prespacer DNA substrates were 5′-radiolabeled with [γ-^32^P]-ATP (Revvity) using the standard Polynucleotide Kinase (Thermo Fisher Scientific) reaction. Only one oligonucleotide of the prespacer was labeled at a time. Afterwards, labeled and unlabeled complementary oligonucleotides were mixed at a 1:1.5 molar ratio and annealed. Unlabeled integration target L-R1-S1-R2-S2 was obtained by PCR using primers CR3fwd and CR3rev. All integration reactions were carried out in a buffer containing 100 mM NaCl, 10 mM Tris-HCl (pH 8.0), 5 mM Mg(CH_3_COO)_2_ and 1 mM DTT. For the kinetic integration assay (Figures 1G and S1E), 100 nM St3 Cas1-Cas2 integrase (C-terminal StrepIITag on St3Cas1) was added to a mixture of 100 nM unlabeled integration target and 100 nM radiolabeled prespacer. The reactions were conducted at 25 °C and stopped at timed intervals (10, 90, 600 and 1800 s) with 20 mM EDTA, followed by heating. Subsequently, the DNA samples were deproteinized with 20 μg of proteinase K (Thermo Fisher Scientific) for 30 min at 50 °C, heated and separated on 10% Urea-PAGE (thermostated at 60 °C). Radioactive bands were visualized by phosphor-imaging using a Typhoon (Cytiva) scanner. End point integration assay in Figure S5C was performed in a similar way. Reactions contained 100 nM St3 Cas1-Cas2 integrase, 100 nM unlabeled integration target, 100 nM radiolabeled prespacer and, if applicable, 100 nM Cas9-RNA or/and 100 nM (Csn2)4 tetramer. Cas9-RNA was preassembled with the radiolabeled prespacer (15 min at 25 °C) prior to adding other proteins and integration target to the mixture. Reactions were carried out for 30 min at 25 °C. For quenching and DNA visualization, the same procedures were implemented as described before.

### Cas9 activity assay

St3Cas9 (WT or Q544A+K706E) cleavage assay was performed for 3 h at 37 °C in a reaction buffer (10 mM Tris-HCl (pH 7.5 at 37 °C), 50 mM NaCl, 10 mM MgCl_2_, 0.1 mg/mL BSA) containing 10 nM Cas9 or 10 nM Cas9:crRNA:tracrRNA mixed with 5 nM pSp1 plasmid^55^ carrying a single DNA target site. To assemble the St3 Cas9:crRNA:tracrRNA complex, crRNA and tracrRNA were pre-annealed (in 33 mM Tris-acetate (pH 7.9 at 37 °C), 66 mM CH_3_COOK, 0.1 mg/mL BSA) and incubated with St3Cas9 in the assembly buffer (10 mM Tris-HCl (pH 7.5 at 25 °C), 100 mM NaCl, 1 mM EDTA, 1 mM DTT) for 15 min at 25 °C prior to the activity assay. Cas9 cleavage reactions were quenched with chloroform and the aqueous phase containing the cleavage products was analyzed by agarose-ethidium bromide electrophoresis.

### Cryo-EM sample preparation

St3Cas9:crRNA:tracrRNA binary complex was reconstituted by mixing purified St3Cas9 with preannealed crRNA:tracrRNA duplex at a 1:1.2 molar ratio in an assembly buffer containing 300 mM NaCl, 30 mM Tris-HCl (pH 7.5), 2 mM Ca(CH_3_COO)_2_ and 1 mM DTT. Incubation time was 10 min at 25 °C. To remove glycerol, sample was dialysed against the assembly buffer overnight and concentrated using an Amicon Ultra protein concentrator (10 kDa MWCO). St3Cas9-RNA ternary complexes with cognate (gs-2015/gs-2016) and non-cognate (gs-2059/gs-2130) DNA were obtained by adding the appropriate DNA to the preassembled Cas9:crRNA:tracrRNA complex and incubating for additional 10 min at 25 °C before dialysis. In the case of St3Cas9 bound to non-cognate DNA, dialysis was performed against a low-salt buffer (100 mM NaCl, 10 mM Tris-HCl (pH 7.5), 2 mM Ca(CH_3_COO)_2_, 1 mM DTT) in order to facilitate DNA binding.

To reconstitute St3 Cas9-Cas1-Cas2-Csn2 supercomplex, Cas9-RNA ternary complex with non-cognate DNA (gs-2059/gs-2130) was assembled and dialysed against the low-salt buffer for 8 h, then St3 Cas1-Cas2 integrase (C-terminal HisTag on St3Cas1) and (Csn2)_4_ tetramer were added to the Cas9-RNA-DNA at a 2:2:1 molar ratio, incubated for 10 min at 4 °C, dialysed overnight and concentrated using an Amicon Ultra concentrator (10 kDa MWCO). Cryo-EM samples of St3 Cas1-Cas2 integrase bound to DNA were prepared by mixing equimolar amounts of St3 Cas1-Cas2 integrase (C-terminal StrepIITag on St3Cas1), prespacer DNA (gs-1754/gs-gs-1755 or gs-1842/gs-1842) and, if applicable, target DNA (gs-2160/gs-2164) in a low-salt buffer (100 mM NaCl, 10 mM Tris-HCl (pH 7.5), 2 mM Ca(CH_3_COO)_2_, 1 mM DTT), followed by dialysis overnight and concentration. St3Csn2^Y212R+I213R+M215R^ complex with Cas1-Cas2 integrase and dsDNA was assembled by mixing together St3Csn2^Y212R+I213R+M215R^, Cas1-Cas2 integrase and 30 bp dsDNA (gs-38/gs-58) at a 2:2:1 molar ratio and incubating for 10 min (25 °C) in the low-salt buffer (100 mM NaCl, 10 mM Tris-HCl (pH 7.5), 2 mM Ca(CH_3_COO)_2_, 1 mM DTT) before dialysing the sample overnight against the same buffer and concentrating using an Amicon Ultra concentrator (10 kDa MWCO).

Control samples with mutant St3Cas9 and St3Csn2 proteins or without dsDNA were prepared as described for the WT St3 supercomplex above, except for the omission of dsDNA (supercomplex formation in the absence of dsDNA tested), or replacement of WT St3Csn2 or St3Cas9 proteins by the mutant variants.

All protein samples were applied on the freshly glow-discharged (20 mA, 45 s in a GloQube Plus device (Quorum Technologies)) Cu 300 mesh R1.2/1.3 holey carbon grids (Quantifoil) and plunge-frozen in liquid ethane using Vitrobot Mark IV (FEI) at 4 °C with a waiting time of 0 s and a blotting time of 5 s under 95% humidity conditions. In the case of St3 Cas1-Cas2 integrase-prespacer complex, the sample was additionally applied on the Cu 300 mesh R1.2/1.3 grid coated with graphene oxide (GO) using a waiting time of 30 s and a blotting time of 4 s under 95% humidity conditions. GO coating was performed as described by Bokori-Brown et al.^56^. Briefly, graphene oxide dispersion (Merck) was diluted in ddH_2_O to 0.2 mg/ml, spun down at 500 g for 60 s to remove aggregates and applied on the carbon side of a freshly glow-discharged (50 mA, 1 min) Cu grid. After incubation of 1 min, the excess of graphene oxide was removed using blotting paper. The grid was washed 3 times with ddH_2_O, dried and subsequently used for plunge-freezing.

### Cryo-EM data collection and image processing

Cryo-EM data collection is summarized in Table S2. Data collection was performed using a Glacios microscope (Thermo Fisher Scientific), operating at 200 kV and equipped with a Falcon 3EC Direct Electron Detector in the electron counting mode (Vilnius University, Vilnius, Lithuania). Data was collected with EPU (v.2.12.1 - v.3.5.1) at a nominal magnification of 92,000×, corresponding to a calibrated pixel size of 1.10 Å, using an exposure of 0.80 e/Å^2^ s^−1^, in 30 frames and a final dose of 30 e/Å^2^, over a defocus range of −1.0 to −2.0 µm. Patch motion correction, CTF estimation, micrograph curation and, in some cases, blob picking with particle extraction were performed in real-time in CryoSPARC Live (v.3.3.1 - v.4.6.0)^57^. Further data processing and final refinement were performed using standard CryoSPARC (v.3.3.1 - v.4.6.0)^57,58^. In all cases, 3DFSC web server^59^ was used to estimate the global resolution at 0.143 FSC cutoff and sphericity values of the final electron density maps, while the local resolution was calculated using CryoSPARC (v.4.6.0)^57^.

#### Typical non-standard CryoSPARC job parameters

Exposure curation: we retained micrographs with the actual defocus range between −0.5 to −2.5 µm, CTF fit resolution exceeding 6 Å, max. in-frame motion below 4 pixels, and total motion below 50 pixels. 2D classification: number of online-EM iterations 30-40, batchsize per class 300-400, number of 2D classes 100-200. Ab-initio reconstruction: initial minibatch size 1000, final minibatch size 1500, number of ab-initio classes 2-3. Non-uniform refinement: enabled ‘optimize per-group CTF params’, ‘Fit Spherical Aberration’, ‘Fit Tetrafoil’ and ‘Fit Anisotropic Mag.’ options. Local refinement: rotation search extent 7 deg., shift search extent 3 Å. 3D classification (without alignment): number of classes 2-3, filter resolution 5-6 Å, simple initialization mode.

#### St3Cas9:crRNA:tracrRNA binary complex

Cryo-EM data for the St3Cas9:crRNA:tracrRNA binary complex was processed as depicted in Figure S9A. Briefly, 860,841 particles were blob-picked and extracted (box size 200 pixels, particle size 90-120 Å) from 1,518 accepted micrographs. After two-dimensional (2D) classification, the selected particles (132,933) were subjected to ab-initio reconstruction into two classes, followed by heterogeneous refinement. Particles from the selected class (108,160) were used for the final reconstruction using non-uniform refinement, local refinement, reference-based motion correction, and the final rounds of non-uniform and local refinement.

#### St3Cas9 ternary complex with cognate DNA

The cryo-EM single-particle reconstruction workflow of St3Cas9 bound to cognate DNA is displayed in Figure S9B. In brief, 894,269 particles of 90-120 Å in size were picked using blob-picker and extracted from 1,657 accepted micrographs using a box size of 200 pixels. After 2D classification, the selected particles (268,073) were subjected to heterogeneous refinement using two volumes obtained from an ab-initio reconstruction. The class with the higher FSC resolution (4.29 Å, 207,195 pct) was further used for non-uniform refinement and local refinement, followed by 3D classification into 2 classes of a similar size. The first 3D class represented St3Cas9 ternary complexes with a partially resolved REC2 domain, while the second 3D class represented a more complete St3Cas9 complex with a partially resolved REC3 domain. The particles from the second 3D class (119,790) were used for the final reconstruction using non-uniform and local refinement.

#### St3Cas9 ternary complexes with non-cognate DNA

Cryo-EM data processing of the St3Cas9 in complex with the non-cognate DNA is represented schematically in Figure S10. To summarize, 3,199,891 particles were picked using a template picker (particle diameter 120 Å) and extracted (200 px) from 3,672 accepted and denoised micrographs, followed by 2D classification. 526,132 particles from the selected 2D classes were used for ab-initio reconstruction and heterogeneous refinement into two classes. Particles from the selected class (380,383 ptc) were subjected to non-uniform refinement and local refinement, followed by 3D classification (5 Å target resolution) into 3 classes using a mask focused onto the DNA-binding groove of the Cas9 PAM-interacting domain. The resultant 3D classes were of similar size and represented different St3Cas9 species as follows: the first class (127,102 ptc) represented the St3Cas9 binary complex, the second class (127,551 ptc) represented St3Cas9 bound to a DNA fragment non-specifically, and the final class (125,730) represented St3Cas9 forming PAM-specific contacts with the DNA. Both DNA-bound St3Cas9 classes were used separately to obtain distinct electron density maps by performing non-uniform and local refinement, reference-based motion correction, and the additional rounds of non-uniform and local refinement. The PAM-bound St3Cas9 particles were subjected to an additional round of 3D classification using a focused mask covering the DNA region. The 3D class (41,668 particles) containing the longest resolved DNA fragment (approx. 25 bp) was locally refined to obtain the final reconstruction for PAM-bound St3Cas9.

#### St3 Cas1-Cas2 integrase bound to prespacer DNA

Cryo-EM processing and 3D reconstruction workflow of the St3 Cas1-Cas2 integrase in complex with the prespacer DNA is visually depicted in Figure S11A. Briefly, 674,498 particles (diameter 100-170 Å) were picked using blob picker and extracted using a box size of 256 pixels from 1,589 accepted micrographs, which were collected from a standard Cu grid. To improve the uniformity of orientation distribution of the integrase-DNA complex, an extra dataset was collected from a graphene-coated (GO) Cu grid, resulting in additional 1,037,877 particles (blob picking, particle size 100-170 Å), which were extracted (box size 256 pixels) from 1,713 accepted and denoised micrographs. After 2D classification, particles from each data set were selected (428,579 ptc and 143,377 ptc from standard and GO grids, respectively) and pooled together to perform an ab-initio reconstruction into two classes, followed by two rounds of heterogeneous refinement in order to separate the full heterohexameric (Cas1)_2_-(Cas2)_2_-(Cas1)_2_ integrases from integrases with an unresolved second Cas1 dimer. The class representing full integrase-DNA complexes (226,248 ptc) was used for two rounds of non-uniform refinement, imposing C2 symmetry constraints. In between refinements the orientation distribution of particles was rebalanced by removing particles from oversampled views (Rebalance Orientations job in CryoSPARC 4.6.0, rebalance percentile 0.90), achieving a final reconstruction from 136,273 particles.

#### St3 Cas1-Cas2 integrase bound to prespacer and target DNA

Cryo-EM data of St3 Cas1-Cas2 integrase bound to prespacer and target DNA was processed as displayed in Figure S11B. Briefly, 1,556,184 particles were blob-picked and extracted (box size 320 pixels, particle size 150-200 Å) from 2,755 accepted micrographs. 2D classification was performed in two rounds, resulting in 129,192 selected particles, which were later subjected to ab-initio reconstruction into two classes and heterogeneous refinement. To obtain the final electron density map, class 2 possessing a higher FSC resolution (5.63 Å, 82,944 pct) was used for non-uniform refinement with applied C2 symmetry, reference-based motion correction, and the additional round of non-uniform refinement with C2 symmetry.

#### Partial St3 Cas1-Cas2 integrase bound to a half-integration product

Reconstruction workflow of the partial St3 Cas1-Cas2 integrase in complex with half-integration DNA product is represented in Figure S12A. In brief, 1,711,840 particles of 120-180 Å in diameter were picked using an automatic blob-based picker and extracted using a box size of 256 pixels from 2,031 accepted micrographs, followed by 2D classification. 345,756 selected particles were subjected to two rounds of heterogeneous refinement using two volumes obtained from an ab-initio reconstruction. Particles from the selected volume class (196,153 ptc) were used for non-uniform refinement, local refinement, and 3D classification (5 Å target resolution, 2 classes) with solvent mask only. The resultant 3D classes represented fully resolved partial (Cas1)_2_-(Cas2)_2_ integrase assemblies with (87,779 ptc) or without (108,374 ptc) poorly resolved density in the position of the second (Cas1)_2_ dimer. For the final reconstruction, the particles from the 3D class (108,374 ptc, no density for the second (Cas1)_2_ dimer) were selected and used for an additional round of local refinement.

#### St3Csn2^Y212R+I213R+M215R^ in complex with Cas1-Cas2 integrase and dsDNA

Cryo-EM data of St3Csn2^Y212R+I213R+M215R^ bound to Cas1-Cas2 integrase and dsDNA was processed as displayed in Figure S12B. In summary, 752,762 particles were picked using a template picker and extracted (box size 256 pixels, particle size 120 Å) from 912 accepted and denoised micrographs. Following 2D classification, selected particles were re-extracted using a box size of 320 pixels, resulting in a total of 245,192 particles that were subjected to two rounds of heterogeneous refinement using volumes generated by an ab-initio reconstruction. Particles from the selected class (92,588 ptc) were used for the final reconstruction via non-uniform refinement and local refinement.

#### St3 supercomplex

The cryo-EM single-particle reconstruction workflow of the St3 supercomplex is shown in Figure 13. Briefly, 6,627 accepted micrographs were denoised, followed by template-based picking and extraction of 2,945,516 particles (diameter 170 Å) using a box size of 336 pixels. After 2D classification, 514,314 selected particles were subjected to two rounds of heterogeneous refinement using volumes obtained from an ab-initio reconstruction. The supercomplex comprised the minor fraction (<25%) of particles in the samples, suggestive of its low stability under our experimental conditions. Particles belonging to the supercomplex (138,824 ptc) were used for non-uniform refinement and local refinement, followed by focused 3D classification (5 Å target resolution) into 2 classes using a mask covering the DNA-binding groove of the Cas9 PAM-interacting domain. The first 3D class (67,204 ptc) represented the supercomplex non-specifically bound to an undistorted DNA fragment (search conformation), and the second 3D class (71,620 ptc) represented the supercomplex forming PAM-specific contacts with the DNA. The PAM-unbound 3D class was subjected to local refinement yielding the consensus map for the supercomplex in the search conformation. The PAM-bound supercomplex class was subjected to further 3D classification using a mask covering the DNA region between PAM and Csn2. The two larger classes (26,925 and 24,394 ptc) represented the PAM-bound supercomplex with bent DNA (the docked conformation) and PAM-bound supercomplex with a well-resolved fragment of straight DNA (the locked conformation). These two classes were subjected to local refinement yielding consensus maps for the supercomplex in the docked and locked conformations.

This was followed by focused refinement of each consensus map using masks covering either Cas9-Cas1 or Csn2-Cas1 subunits of the supercomplex. A composite map for each supercomplex conformation was obtained by combining the consensus map with two focused refinement maps using phenix.combine_focused_maps function in Phenix (v. 1.21.2-5419) with the ‘rigid_body_refinement_single_unit’ setting set to ‘True’ ^60^.

For the control samples obtained without dsDNA or with mutant St3Cas9 or St3Csn2 proteins, smaller data sets (less than 1000 micrographs) were collected, and analyzed as above; obtained 2d class averages were used to assess the presence/absence of supercomplex particles.

### Cryo-EM model building, refinement and analysis

Initial models (either experimental structures or Alphafold 2^61^ models, Table S2) were fitted into the cryo-EM maps using ChimeraX (v.1.7)^62^. Protein rebuilding and manual building of DNA and crRNA:tracrRNA were performed using Coot (v.0.9.8.1)^63^. Model refinement was performed using phenix.real_space_refine (v. 1.21.2-5419)^60^, refinement statistics are summarized in Table S2. Structure overlays and generation of structural images were performed using ChimeraX (v.1.7)^62^; protein interface surfaces were analyzed using VoroContacts web server^64^ with a 4.0 Å distance cut-off; sequence alignments were generated using Clustal Omega (1.2.4)^65^ and ESPript 3.0^66^.

#### Model building of St3Cas9 complexes

The initial protein model for St3Cas9 was generated using AlphaFold 2^61^. First, separate domains of the St3Cas9 model were rigid-body fitted into the cryo-EM map of St3Cas9 ternary complex with PAM-containing non-cognate DNA, followed by protein rebuilding and manual building of DNA and crRNA:tracrRNA. The final model of the St3 ternary complex bound to PAM-containing non-cognate DNA (PDB: 9h2g) covers protein residues 3-258, 265-765, 777-1385, crRNA residues -11 to 14, tracrRNA residues 8-72, and has a PAM-bound DNA duplex of 24 bp. This model without DNA was further used as the initial model for St3Cas9:crRNA:tracrRNA binary complex (PDB: 9h21) and St3Cas9 bound to non-cognate DNA with no interactions with the PAM region (PDB: 9h2m). Though in the latter case low resolution of the cryo-EM map in the DNA region did not allow us to trace the DNA sequence, we have manually inserted an undistorted B-form DNA generated in Coot with a sequence matching the central part of the oligoduplex used in sample preparation. The final models of interest were obtained after several rounds of manual adjustment in Coot and real-space refinement in Phenix. The model of St3Cas9:crRNA:tracrRNA binary complex (PDB: 9h21) covers protein residues 3-249, 286-765 and 777-1385, crRNA residues -10 to 11, and tracrRNA residues 11-72. The model of St3Cas9 ternary complex bound to a non-specific DNA fragment (PDB: 9h2m) covers the same protein and RNA residues as the initial model (PDB: 9h2g) and also contains a 27 bp DNA duplex. In the case of St3Cas9 ternary complex with cognate-DNA, Alphafold 2 model was fitted into the cryo-EM map with minor adjustments, followed by manual building of DNA, crRNA:tracrRNA, and crRNA:DNA heteroduplex. This model (PDB: 8pj9) covers around 76% of the protein sequence (residues 3-175, 308-765, 935-1018, 1040-1049, 1060-1385), contains crRNA -20 to 15 and tracrRNA 7-72 nucleotides, and a DNA molecule (target strand: -9 to 20 residues, non-target strand: 0-9 residues) forming a heteroduplex with crRNA upstream of the PAM sequence. The obtained map allowed us to build only one nucleotide of the non-target strand adjacent to the PAM.

#### Model building of St3 Cas1-Cas2 integrase complexes

The initial protein model for St3 Cas1-Cas2 integrase was generated using AlphaFold 2^61^. This model was used to build the atomic models of integrase-prespacer (PDB: 9h1h) and integrase-prespacer-target (PDB: 9h1v) complexes performing similar procedures as mentioned above. DNA was built manually. The final model of St3 Cas1-Cas2 integrase in complex with prespacer DNA (PDB: 9h1h) covers 3-289 and 3-288 residues of the catalytic Cas1* and the non-catalytic Cas1 subunits, respectively, together with 5-111 amino acids of the Cas2 subunits, and contains two DNA molecules: a 22 bp prespacer with 4-nt 3′-overhangs at each end and a DNA fragment bound non-specifically on the target DNA-binding side of the integrase. Due to the low cryo-EM map resolution of the non-specifically bound DNA we could not trace the DNA sequence, thus a 12 bp fragment of undistorted B-form DNA generated in Coot with the sequence corresponding to the central part of the oligoduplex used for sample preparation was inserted into the model.

The final model of St3 Cas1-Cas2 integrase bound to prespacer and target DNA (PDB: 9h1v) covers 1-289 residues of the catalytic Cas1* subunits, 2-290 residues of the non-catalytic Cas1 subunits, 4-111 residues of the Cas2 subunits, and has a 22 bp prespacer with 4-nt 3′-overhangs as in the previous model (PDB: 9h1h) together with a bent 48 bp target DNA molecule. The (Cas1)_2_-(Cas2)_2_ assembly with DNA from this model (PDB: 9h1v) was used as the initial model for building of the partial St3 Cas1-Cas2 integrase complex bound to the half-integration product (PDB: 9h6t). Protein residues of this final model are as follows: 4-289 residues of the catalytic Cas1* subunit, 2-289 residues of the non-catalytic Cas1 subunit, 5-110 residues of the Cas1-proximal Cas2 subunit, and 6-100 residues of the Cas1-distal Cas2 subunit. The half-integration DNA product consists of a 21 bp prespacer with a 4-nt 3′-overhang integrated at the leader-repeat junction of a target DNA molecule, comprising 20 bp Repeat and 6 bp Leader sequences.

(Cas1)_2_ dimer from the PDB: 9h1v St3 Cas1-Cas2 model together with AlphaFold 2^61^ model of Csn2 were used as initial models to build the atomic model of St3Csn2^Y212R+I213R+M215R^ complex with dsDNA and Cas1-Cas2 integrase interacting via the alternative Cas1-Csn2 interaction surface (PDB: 9h72). The map allowed us to resolve 2-289 residues of the catalytic Cas1* subunit, 3-290 residues of the non-catalytic Cas1 subunit, a fragment of the Cas2 subunit (100-111 residues), 1-218 residues of all four Csn2 chains, and a DNA fragment encircled by the homotetrameric Csn2 ring. Due to the low cryo-EM map resolution of the non-specifically bound DNA we could not trace the DNA sequence, thus a 12 bp fragment of undistorted B-form DNA generated in Coot with the sequence corresponding to the oligoduplex used for sample preparation was inserted into the model.

#### Model building of the St3 supercomplexes

AlphaFold 2^61^ model of St3Csn2, Cas1 dimer bound to a Cas2 fragment from the St3 integrase structure (PDB: 9h1v), and St3Cas9 ternary complex bound to non-cognate DNA were used as initial models to build the atomic models of St3 Cas9-Cas1-Cas2-Csn2 supercomplexes. The PAM-bound supercomplexes were built using the model of Cas9 specifically bound to PAM in the PAM-containing non-cognate DNA (PDB: 9h2g), while the supercomplex forming only non-specific interactions with the DNA fragment was built using an analogous model of St3Cas9 (PDB: 9h2m).

The final model of the PAM-bound St3 supercomplex in the docked conformation (PDB: 9hp8) covers 3-765, and 777-1385 Cas9 residues, -10 to 18 crRNA residues, 4-72 tracrRNA residues, 2-289 residues of both the catalytic Cas1* and the non-catalytic Cas1 subunits, contains a fragment of the Cas2 subunit (101-110 residues), has two fully resolved Cas9-proximal Csn2 chains (chain O: 1-48, 52-218 residues, chain L: 1-219 residues) and partially resolved two distal Csn2 chains (chain N: 18-38, 58-197 residues, chain M: 16-39, 58-197 residues), and includes a 43 bp DNA duplex. Only in the PAM-surrounding region the map quality enabled de-novo modeling of the DNA; the remaining part was built in Coot as an undistorted B-form DNA duplex with the sequence matching that of the oligoduplex used in the sample, and was fitted into the low-resolution part of the map between Cas9 and Csn2 using the intra-molecular restraints functionality in Coot^63^.

The final model of the PAM-bound St3 supercomplex in the locked conformation (PDB: 9q85) covers 3-195, 205-229, 233-765, and 777-1385 Cas9 residues, -10 to 18 crRNA residues, 4-72 tracrRNA residues, 2-289 residues of both the catalytic Cas1* and the non-catalytic Cas1 subunits, contains a fragment of the Cas2 subunit (101-110 residues), has two fully resolved Cas9-proximal Csn2 chains (chain O: 1-218 residues, chain L: 1-219 residues) and partially resolved two distal Csn2 chains (chain N: 18-38, 58-197 residues, chain M: 16-39, 58-197 residues), and includes a 43 bp DNA duplex. The DNA region adjacent to PAM was built in Coot de novo manually, the remaining part was built as an undistorted B-form DNA duplex with the sequence matching that of the oligoduplex used in the sample.

The final model of the PAM-unbound St3 supercomplex (the search conformation, PDB: 9hp9) covers 3-195, 205-229, 233-257, 264-765, and 777-1385 Cas9 residues, -10 to 18 crRNA residues, 4-72 tracrRNA residues, 2-289 residues of both the catalytic Cas1* and the non-catalytic Cas1 subunits, contains a fragment of the Cas2 subunit (101-110 residues), has two fully resolved Cas9-proximal Csn2 chains (chain O: 1-218 residues, chain L: 1-219 residues) and partially resolved two distal Csn2 chains (chain N: 18-38, 58-197 residues, chain M: 16-39, 58-197 residues), and includes a 43 bp DNA fragment. The DNA was generated in Coot as an undistorted B-form DNA using the sequence of the oligoduplex present in the sample, and was inserted into the low resolution cryo-EM map of the DNA in the same orientation as in the docked conformation supercomplex described above.

### Protein stability measurements

The stability of St3Cas9 and St3Csn2 wt and mutant variants was assessed by performing nano differential scanning fluorimetry (nanoDSF) with a Prometheus NT.48 instrument (NanoTemper Technologies), equipped with backreflection module. Protein samples were prepared in a buffer containing 500 mM NaCl, 50 mM Tris-HCl (pH 8.0), 2 mM Ca(CH₃COO)₂, and 1 mM DTT. The concentrations were 0.2 mg/mL for St3Cas9 and ∼1.3 mg/mL for St3Csn2. Protein samples were loaded into standard glass capillaries and subjected to thermal stress from 20 °C to 90 °C at a rate of 1 °C/min. The fluorescence intensity ratio (F350/F330) was plotted against the temperature, and the melting temperature (T_m_) was determined from the maximum of the first derivative using PR.ThermControl V2.1 software. Protein aggregation was assessed simultaneously using backreflection optics, and the aggregation onset temperature (T_agg_) was determined with the same software. For all proteins, the mean and standard deviation of T_m_ and T_agg_ were calculated from triplicate measurements.

### SEC-MALS

The molecular weight of the studied complexes was determined using size-exclusion chromatography coupled with multi-angle laser light scattering (SEC-MALS) on an HPLC system (Waters Breeze) with a miniDAWN MALS detector (Wyatt) and an Optilab refractive index detector (Wyatt). St3 Cas1-Cas2 integrase–prespacer–target, St3Cas9-RNA, and the supercomplex samples were prepared as described above for the respective cryo-EM samples. In the case of St3Cas9-RNA, the dialysis step was omitted. The St3Csn2 SEC-MALS sample was prepared by diluting the protein to a final concentration of 5 µM (Csn2)₄ using a buffer containing 200 mM NaCl and 20 mM Tris-HCl (pH 8.0). A total of 200 µL of protein samples was loaded onto a Superose 6 Increase 10/300 GL column (Cytiva) equilibrated with the appropriate SEC-MALS buffer. Data acquisition and molecular weight determination were performed using ASTRA 7 software (Wyatt) using the refractive index signal for the assessment of sample concentration.

## Data availability

All data are available in the manuscript and the supplementary material. The EM densities and model coordinates have been deposited in the Electron Microscopy Data Bank and Protein Data Bank under the accession codes EMD-51787/PDB:9h21 (St3Cas9 binary complex), EMD-17702/PDB:8pj9 (St3Cas9 ternary complex with cognate DNA), EMD-51812/PDB:9h2m (St3Cas9 ternary complex with non-cognate DNA/unbound PAM), EMD-51801/PDB:9h2g (St3Cas9 ternary complex with non-cognate DNA/bound PAM), EMD-51766/PDB:9h1h (St3 Cas1-Cas2 integrase bound to prespacer DNA), EMD-51773/PDB:9h1v (St3 Cas1-Cas2 integrase bound to prespacer and target DNA), EMD-51900/PDB:9h6t (asymmetric St3 Cas1-Cas2 integrase bound to a half-integration product), EMD-51909/PDB:9h72 (Csn2^Y212R+I213R+M215R^ bound to Cas1-Cas2 integrase via Interface 4). The EM densities of the supercomplexes (composite map, consensus map, focused map 1 obtained with Cas1-Cas9 mask and focused map 2 obtained with Cas1-Csn2 mask) have been deposited in the Electron Microscopy Data Bank under the accession codes EMD-52324/ EMD-52321/ EMD-52322/ EMD-52323 (search conformation), EMD-52320/ EMD-52317/ EMD-52318/ EMD-52319 (PAM-bound, docked conformation), and EMD-52886/EMD-52883/EMD-52884/EMD-52885 (PAM-bound, locked conformation). The supercomplex model coordinates have been deposited in the Protein Data Bank under the accession codes 9hp9 (search conformation), 9hp8 (PAM-bound, docked conformation), and 9q85 (PAM-bound, locked conformation).

## ACKNOWLEDGEMENTS

We thank Rimantas Sapranauskas and Gediminas Drabavicius for St3 protein expression plasmids, Philippe Horvath for a kind gift of *S. thermophilus* DGCC 7710 strain and D2972 bacteriophage, Martin Wilkinson for the help with design of Interface 1 and Interface 4 mutations, and Miglė Kazlauskienė for discussions and critical reading of the manuscript. This work was funded by the Research Council of Lithuania (grant S-MIP-19-32 to GS).

## AUTHOR CONTRIBUTIONS

Conceptualization: UG, GG, VS, GS

Funding acquisition: GS

Methodology: UG, AS, GS

Formal analysis: UG, GS

Investigation: UG, AS, GT, GS

Resources: UG, GG, AS, GS

Visualization: GS, UG

Project administration: GS

Supervision: GS

Writing - original draft: GS, UG, VS.

## Competing interests

GG is an employee, board member, and has financial interests in Caszyme. VS is a chairman and has financial interests in Caszyme.

## SUPPLEMENTAL FIGURES

**Figure S1.**
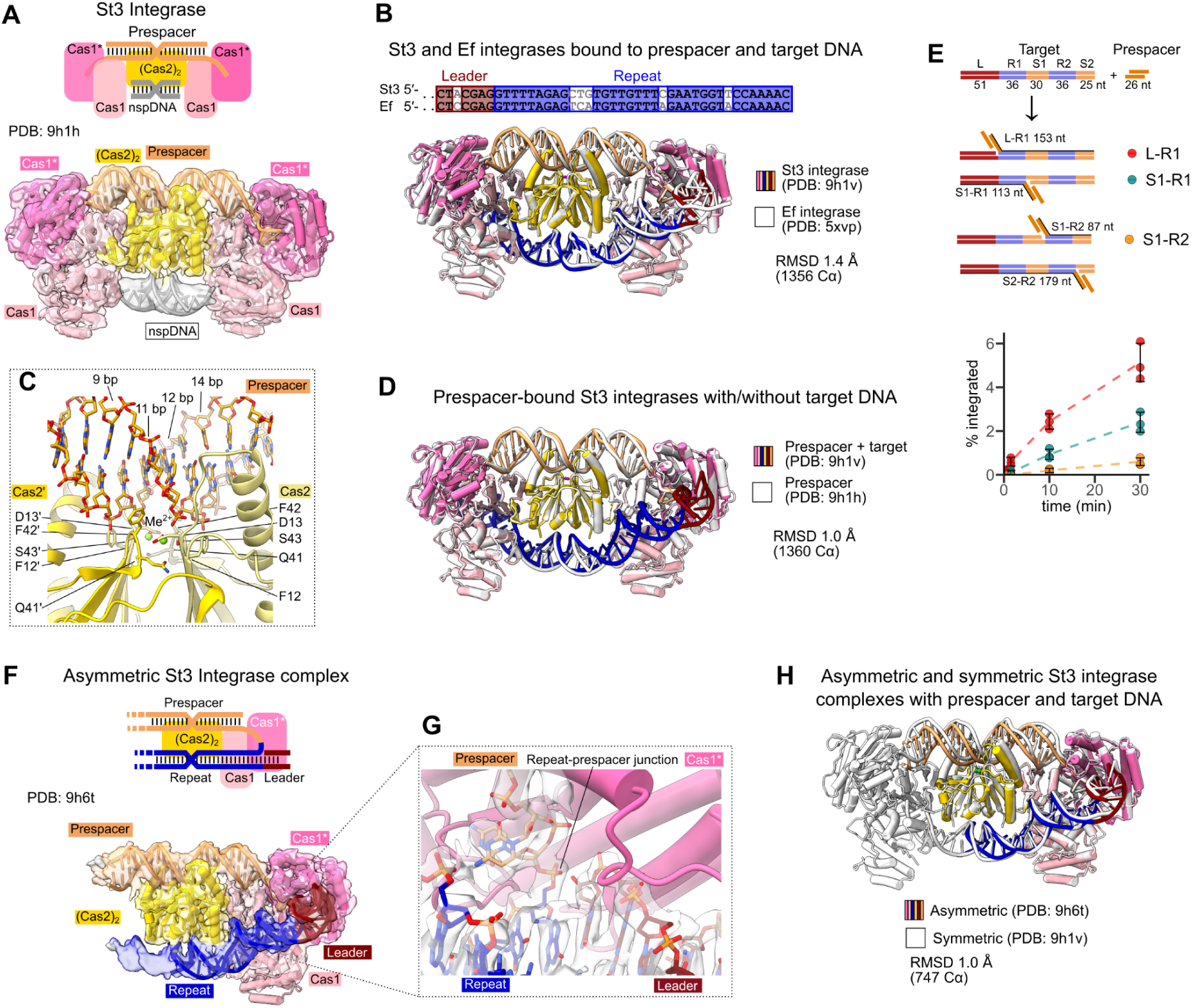
Structural features of the St3 integrase complex. (A) Structure of St3 Cas1-Cas2 integrase complex bound to prespacer DNA. Top: schematic representation of the (Cas1)_2_-(Cas2)_2_-(Cas1)_2_ complex with prespacer DNA (22 bp duplex region with 4 nt 3′-overhangs) and a non-specific DNA fragment (nspDNA, the second copy of prespacer DNA) bound at the integration target binding site; bottom: unsharpened cryo-EM map and atomic model. (B) Comparison of St3 and *E. faecalis* (Ef) Cas1-Cas2 integrases. Top: alignment of St3 and Ef Leader (red) and Repeat (blue) sequences. Identical nucleotides are in colored blocks. Bottom: overlay of St3 and Ef Cas1-Cas2 integrase complexes bound to prespacer and integration target (Leader-Repeat) DNA. St3 Cas1-Cas2 integrase is shown in color, Ef Cas1-Cas2 integrase is shown in white. Sequence identity/similarity for the St3 and Ef integrase proteins is 62/77 % for Cas1, and 70/87 % for Cas2. (C) Divalent cation-mediated contacts of the integrase complex to the central part (11-12 bp) of the 22 bp prespacer fragment (samples were prepared in Ca^2+^-containing buffer). (D) Overlay of St3 Cas1-Cas2 integrase complexes bound to either both prespacer and integration target DNA (shown in color) or prespacer DNA only (shown in white; note that the target-DNA binding site close to Cas2 subunits is occupied by a fragment of non-specifically bound DNA). (E) Time course of St3 integrase-catalyzed prespacer integration reaction. The reactions contained 100 nM integration target, 100 nM radiolabeled prespacer and 100 nM St3 integrase. The ‘S2-R2’ product under our experimental conditions was not detectable (estimated amount < 0.02%). Error bars denote standard deviation of 3 independent experiments. (F) Structure of the asymmetric St3 Cas1-Cas2 integrase complex with unresolved second Cas1 dimer bound to a half-integration product. Top: schematic representation of the (Cas2)_2_-(Cas1)_2_ integrase subcomplex resolved in the cryo-EM map bound to the half-integration product; bottom: unsharpened cryo-EM map and the atomic model. Cas1* denotes the catalytic Cas1 subunit within the Cas1 homodimer. (G) Sharpened cryo-EM map of the asymmetric St3 integrase complex in the vicinity of the repeat-prespacer junction. (H) Overlay of the asymmetric St3 Cas1-Cas2 integrase complex bound to a half-integration product (shown in color) and the symmetric (full) St3 Cas1-Cas2 integrase complex bound to prespacer and target DNA (shown in white).

**Figure S2.**
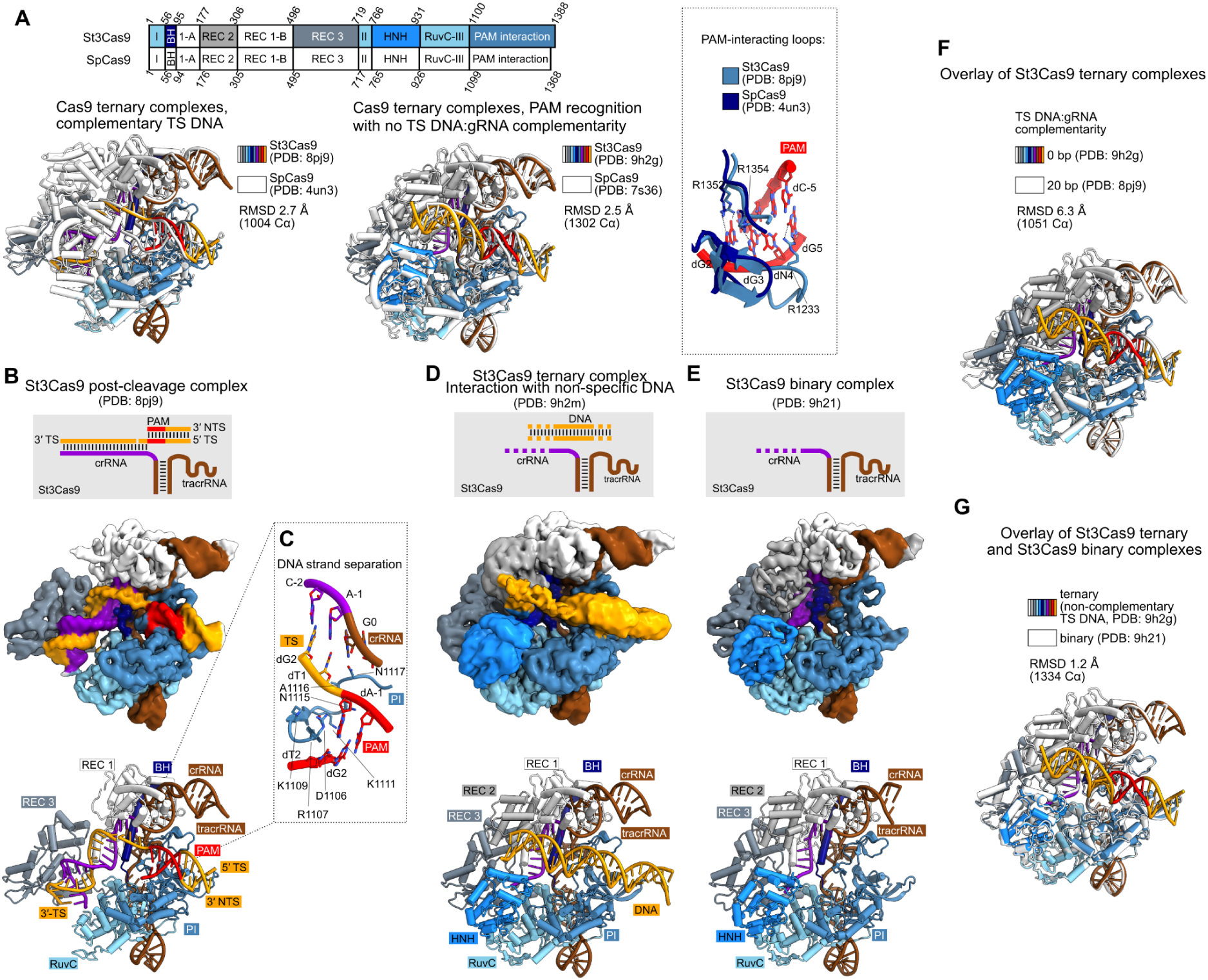
Structural features of St3Cas9. (A) Comparison of St3Cas9 with SpCas9 (sequence identity 57.4 %, similarity 73.4 %). Top: domain organization of St3Cas9 and SpCas9; bottom left: overlay of ternary Cas9 complexes formed with cognate DNA (full TS DNA:guideRNA complementarity); bottom right: overlay of ternary Cas9 complexes recognizing PAM sequences within non-complementary DNA. St3Cas9 elements are shown in color, SpCas9 is shown in white. An overlay of PAM recognition elements of St3Cas9 (PAM: 5′-NGGNG-3′) and SpCas9 (PAM: 5′-NGG-3′) is shown on the right. PAM DNA is shown only for St3Cas9. The St3Cas9 loop that contacts the 5th G (residues 1220-1238) is shorter in SpCas9 (residues 1211-1222). (B) Structure of the St3Cas9:crRNA:tracrRNA complex with a cognate DNA cleavage product. Top: schematic representation of the complex; center: unsharpened cryo-EM map; bottom: atomic model. Structural elements of St3Cas9 are colored as in panel (a). (C) DNA strand separation in the St3Cas9 post-cleavage complex. (D) Structure of the St3Cas9:crRNA:tracrRNA complex interacting with a non-specific DNA fragment. Top: schematic representation of the complex; center: unsharpened cryo-EM map; bottom: atomic model. (E) Structure of the St3Cas9:crRNA:tracrRNA binary complex. Top: schematic representation of the complex; center: unsharpened cryo-EM map; bottom: atomic model. (F) and (G) Overlays of St3Cas9 ternary complexes (complementary/non-complementary DNA with bound PAM) with the St3Cas9 binary complex. Structural elements of St3Cas9 in all panels are colored according to the scheme in panel (a).

**Figure S3.**
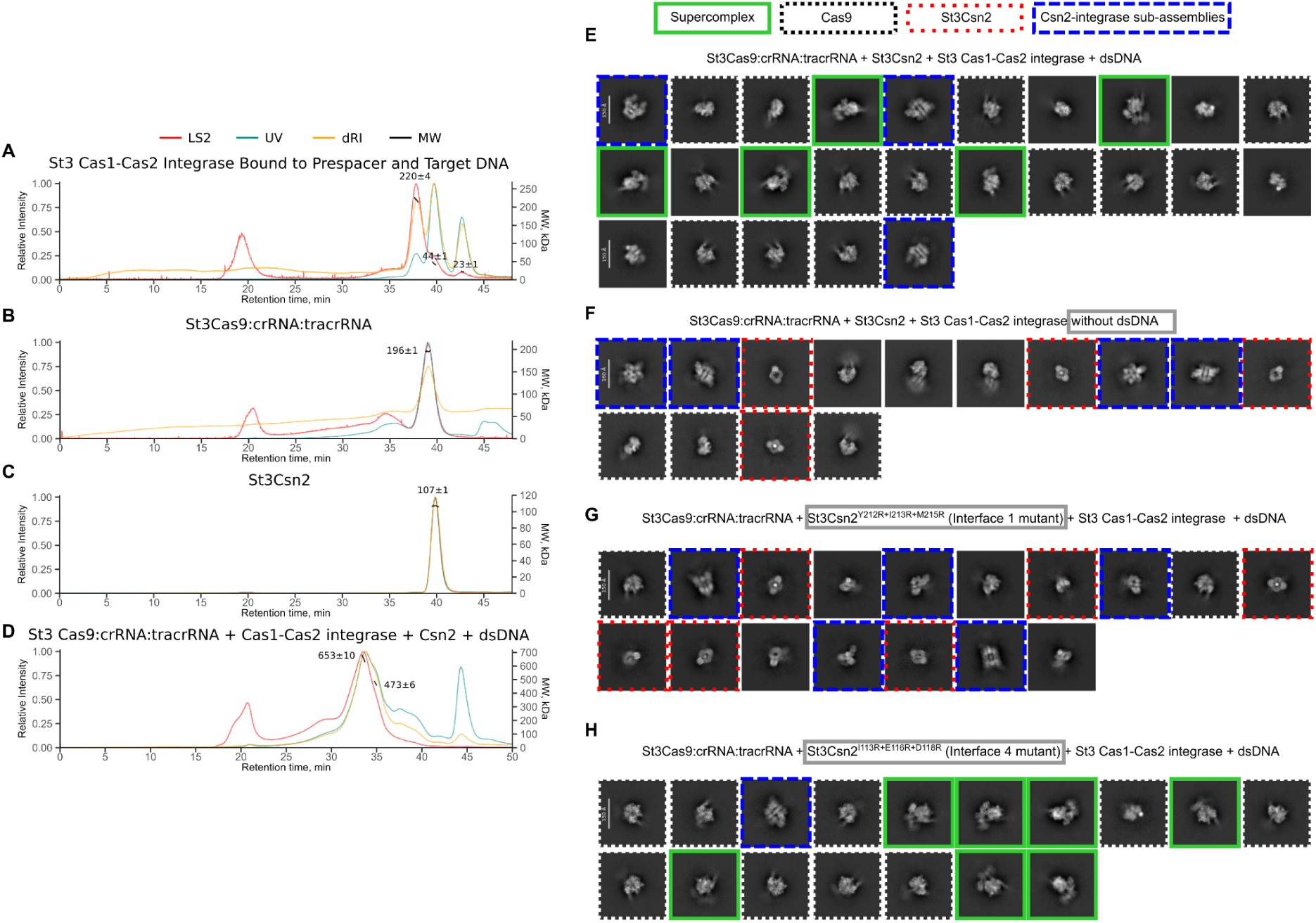
SEC-MALS analysis of St3 proteins and formation of the St3 supercomplex *in vitro*. (A) SEC-MALS analysis of St3 Cas1-Cas2 integrase with prespacer and target DNA. Theoretical MW of St3 (Cas1)_2_-(Cas2)_2_-(Cas1)_2_ integrase complex bound to 1 copy of each DNA is 213 kDa. (B) SEC-MALS analysis of St3Cas9:crRNA:tracrRNA complex, theoretical MW 200 kDa. (C) SEC-MALS analysis of St3Csn2, theoretical MW for the homotetramer is 110 kDa. (D) SEC-MALS analysis of a mixture of all system components with dsDNA; theoretical MW for the 1:1:1:1 Cas9-RNA:(Csn2)_4_:(Cas1)_2_-(Cas2)_2_-(Cas1)_2_ integrase:dsDNA complex is 508 kDa. In panels (A)-(D) the relative light scattering (LS2), absorption at 280 nm (UV), and refractive index (RI) signals are shown. MW values calculated for the respective peaks are indicated. (E)-(H) Cryo-EM analysis of St3 protein samples. All samples were prepared and analyzed as described for the Methods section for the standard WT supercomplex (shown in (E)), except sample in (F) lacked dsDNA, sample in (G) was prepared with the Interface 1 St3Csn2 mutant (St3Csn2^Y212R+I213R+M215R^), whereas sample in panel (H) was prepared with the Interface 4 St3Csn2 mutant (St3Csn2^I113R+E116R+D118R^). The representative high resolution 2D class averages for the respective samples are shown. The identifiable 2D classes corresponding to the supercomplex, Cas9-RNA, Csn2 and Csn2-integrase subcomplexes are marked as shown at the top of (E).

**Figure S4.**
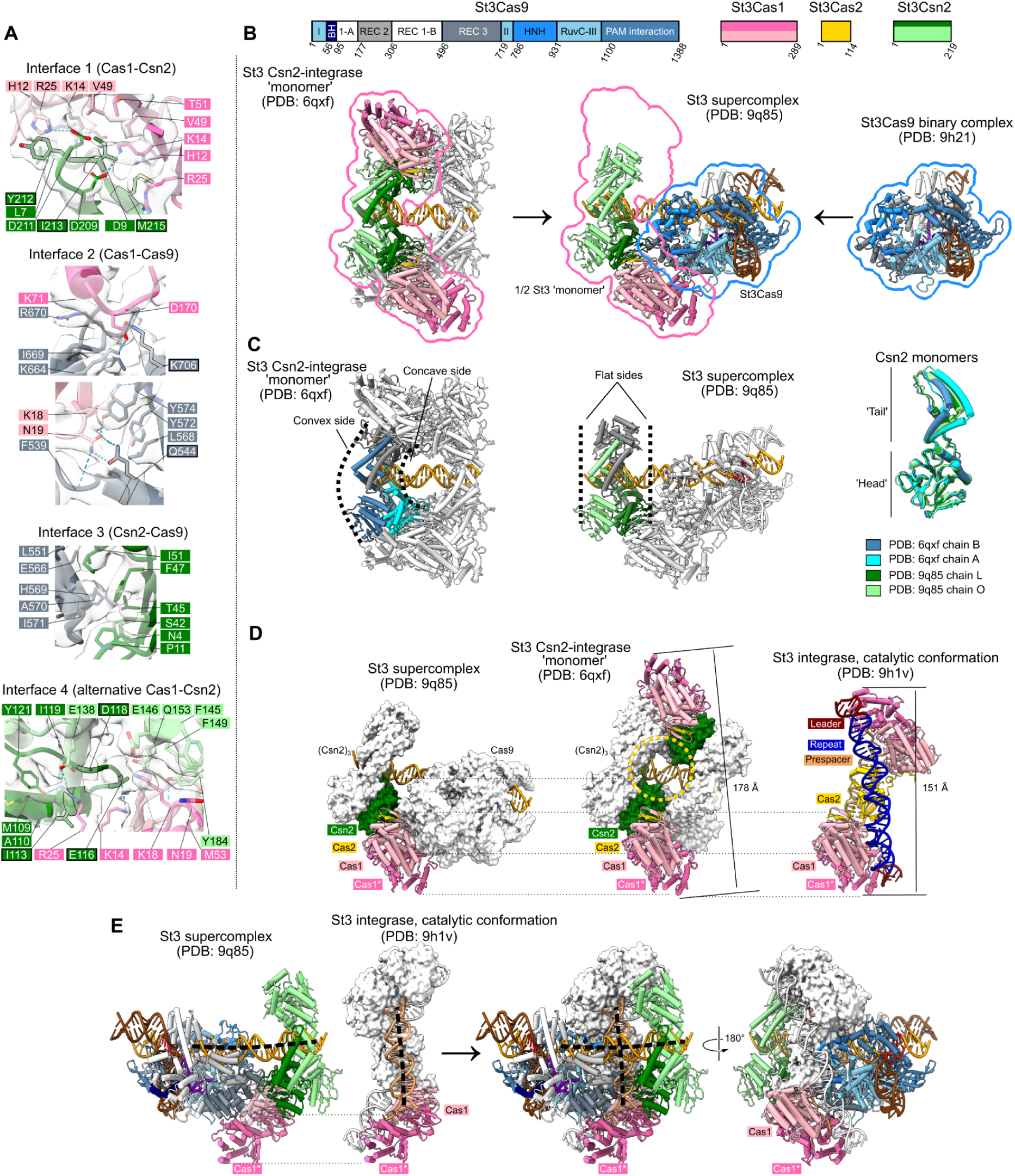
Overall architecture of the St3 supercomplex. (A) Zoom-in views of Cas1-Csn2, Cas9-Cas1 and Cas9-Csn2 interface regions. Residues subjected to mutagenesis are highlighted by black outlines. Sharpened cryo-EM map (EMD-52324, contour level 4) is shown for the indicated residues. (B) Top: domain organization and coloring scheme of the St3 system components. Bottom: the supercomplex (center) corresponds to one half of the St3 Csn2-integrase ‘monomer’ assembly (PDB: 6qxf, left), which includes a single Csn2 tetramer and one Cas1-Cas2 integrase (encircled in pink, only one Cas1 dimer resolved in the supercomplex structures) bound to St3Cas9 (right, encircled in blue). The half of the ‘monomer’ assembly replaced by St3Cas9 is shown in white. (C) Comparison of homotetrameric Csn2 ring conformation in the St3 ‘monomer’ complex (left) and the St3 supercomplex (center). Equivalent Csn2 rings are shown in color, other protein subunits are shown in white. Right: different orientation of the Csn2 ‘tail’ subdomain relative to the ‘head’ in the ‘monomer’ assembly results in a Csn2 ring with concave/convex sides (left); Csn2 ‘tails’ in all Csn2 subunits of the supercomplex adopt intermediate position relative to the ‘head’ subdomain, resulting in a symmetric Csn2 ring with flat sides (center). (D) Arrangement of Cas1 and Cas2 subunits in the St3 supercomplex (left), St3 Csn2-integrase ‘monomer’ complex (center), and the catalytic integrase conformation (right). The structures are aligned based on superposition of Cas1-Cas2 subunits (bottom of the structures, shown as pink cartoons). The Cas1-contacting Csn2 subunits in the super- and ‘monomer’- complexes are shown as green surfaces, the remaining Csn2 subunits and Cas9 are shown as white surfaces. The presumed position of Cas2 dimer within the ‘monomer’ complex is marked by a yellow circle. The St3 integrase is significantly longer in the ‘monomer’ complex than in the catalytic conformation (the distances shown are between His138 residues of the catalytic subunits Cas1*). (E) Structures of St3 integrase in the catalytic conformation (white surface) and the St3 supercomplex are aligned based on superposition of Cas1 subunits (bottom of the structures, shown as pink cartoons). Cas1-Cas2 integrase in the catalytic conformation would clash with Cas9, whereas the DNA in the supercomplex and the prespacer bound to the integrase (black dashed lines) form an approximately right angle.

**Figure S5.**
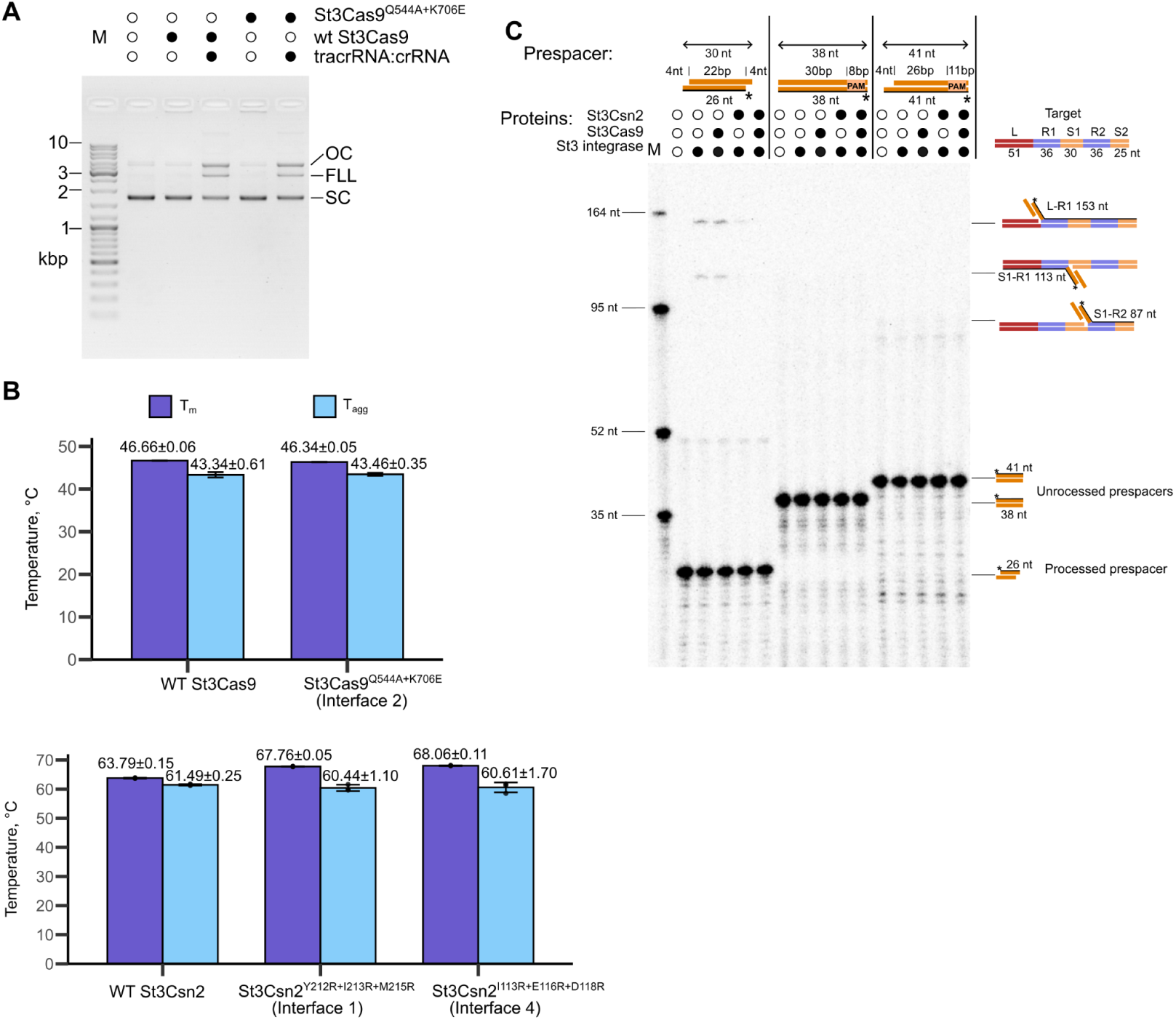
Characterization of mutant St3 proteins and requirements for St3 integrase reactions *in vitro*. (A) Activity of St3Cas9 with mutated Cas9-Cas1 interface (Interface 2) residues. The reactions contained 5 nM pSp1 plasmid (a pUC18 derivative containing a single cognate sequence^55^), and 10 nM St3Cas9 (WT or double mutant St3Cas9^Q544A+K706E^), either with or without 10 nM crRNA:tracrRNA specific for the target site in pSp1 plasmid. Incubation time was 3 h at 37 °C in a buffer containing 50 mM NaCl, 10 mM MgCl_2_, 10 mM Tris-HCl (pH 7.5 at 37 °C) and 0.1 mg/mL BSA. Lane ‘M’ - DNA size marker; ‘OC’ - open circular (nicked) plasmid DNA; ‘FLL’ - full-length linear (cleaved) plasmid, ‘SC’ - supercoiled (intact) plasmid. (B) WT and mutant St3Cas9 and St3Csn2 protein stability measurements. The melting temperature (T_m_), calculated based on the ratio of fluorescence intensity at 350 nm and 330 nm, and the aggregation onset temperature (T_agg_), calculated based on the dynamic light scattering signal, are reported. Results of at least 3 independent measurements (mean value ± stdev) are shown. (C) Activity of St3 Cas1-Cas2 integrase in the presence of St3Csn2 and St3Cas9 and different prespacer DNA variants. The experimental setup was as in Figure 1G, except the reactions were supplemented with St3Cas9:crRNA:tracrRNA and/or St3Csn2, or different variants of radiolabeled prespacer DNA. Integrase activity is observed only with the pre-processed prespacer.

**Figure S6.**
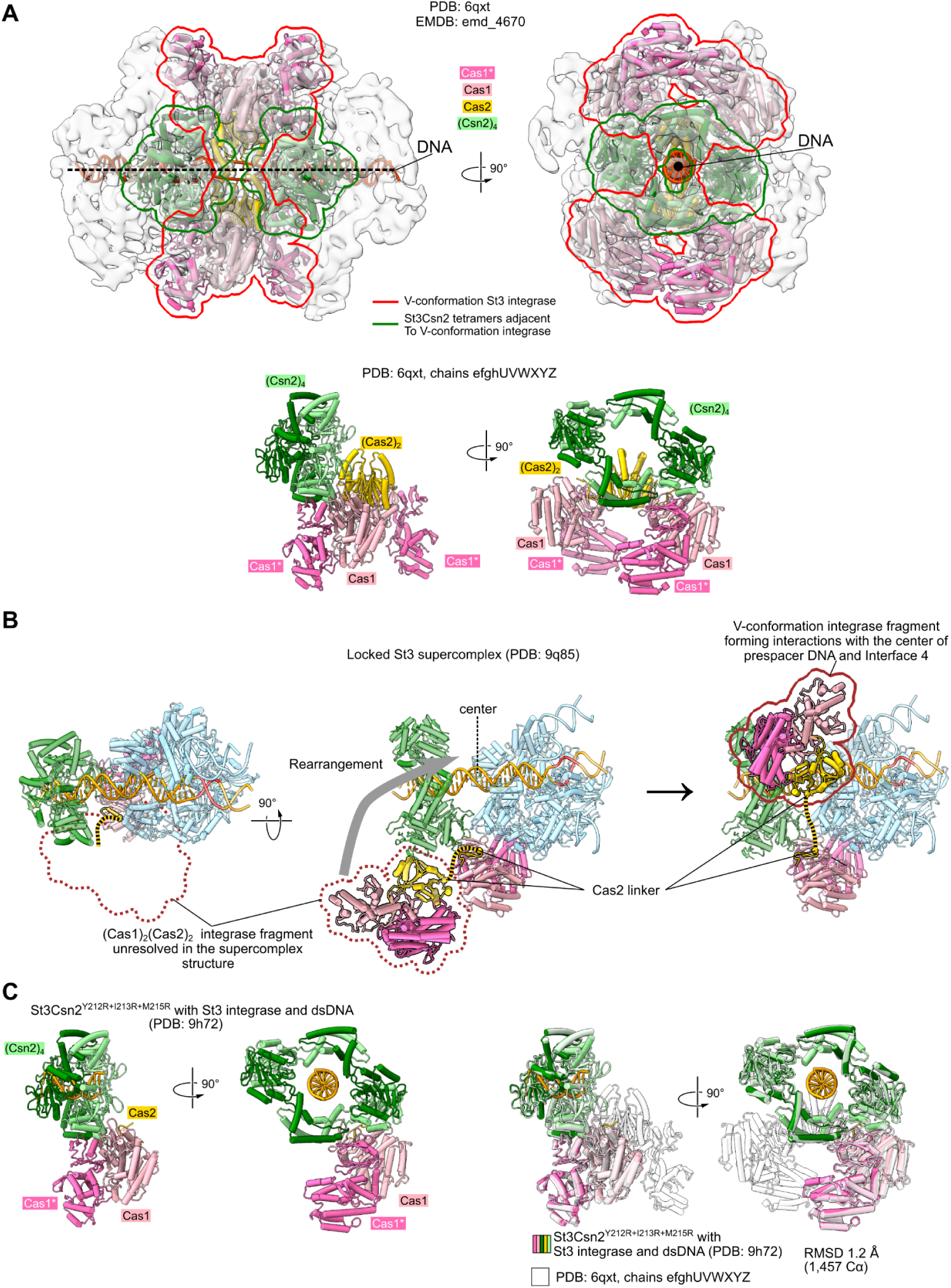
Interface 4 in the experimentally observed Csn2 - integrase assemblies. (A) St3 Csn2-Cas1-Cas2 ‘dimer’ assembly, PDB: 6qxt^30^. The V-conformation integrase complexes forming the central part of the ‘dimer’ assembly (PDB: 6qxt chains U,V,W,X,Y,Z and u,v,w,x,y,z) are encircled in red; Csn2 tetramers that form Interface 4 interactions with the V-conformation integrases (PDB: 6qxt chains e,f,g,h and E,F,G,H) are encircled in green. The Interface V-conformation integrase/Csn2 tetramer sub-assembly (PDB: 6qxt chains U,V,W,X,Y,Z,e,f,g,h) is shown below the ‘dimer’ assembly. (B) Putative rearrangement of integrase fragment, which is unresolved in the supercomplex structure (Cas1 dimer and most of the Cas2 dimer, indicated by a dotted brown line), into a position (solid brown outline) that enables the formation of Interface 4 between Cas1 and Csn2, and Cas2-DNA interactions at the center of the captured prespacer. The 10 aa linker (yellow/black stripes) connects the G90 residue of the V-conformation integrase fragment (Cas1)_2_(Cas2)_2_ docked on the supercomplex, to the E101 residue of the Cas2 C-terminal tail that is resolved in the supercomplex structure. (C) Comparison of the Interface 4-forming St3Csn2^Y212R+I213R+M215R^ complex with St3 integrase and dsDNA (shown in color) and the Interface 4-forming V-conformation integrase/Csn2 tetramer sub-assembly (PDB: 6qxt chains U,V,W,X,Y,Z,e,f,g,h, shown in white).

**Figure S7.**
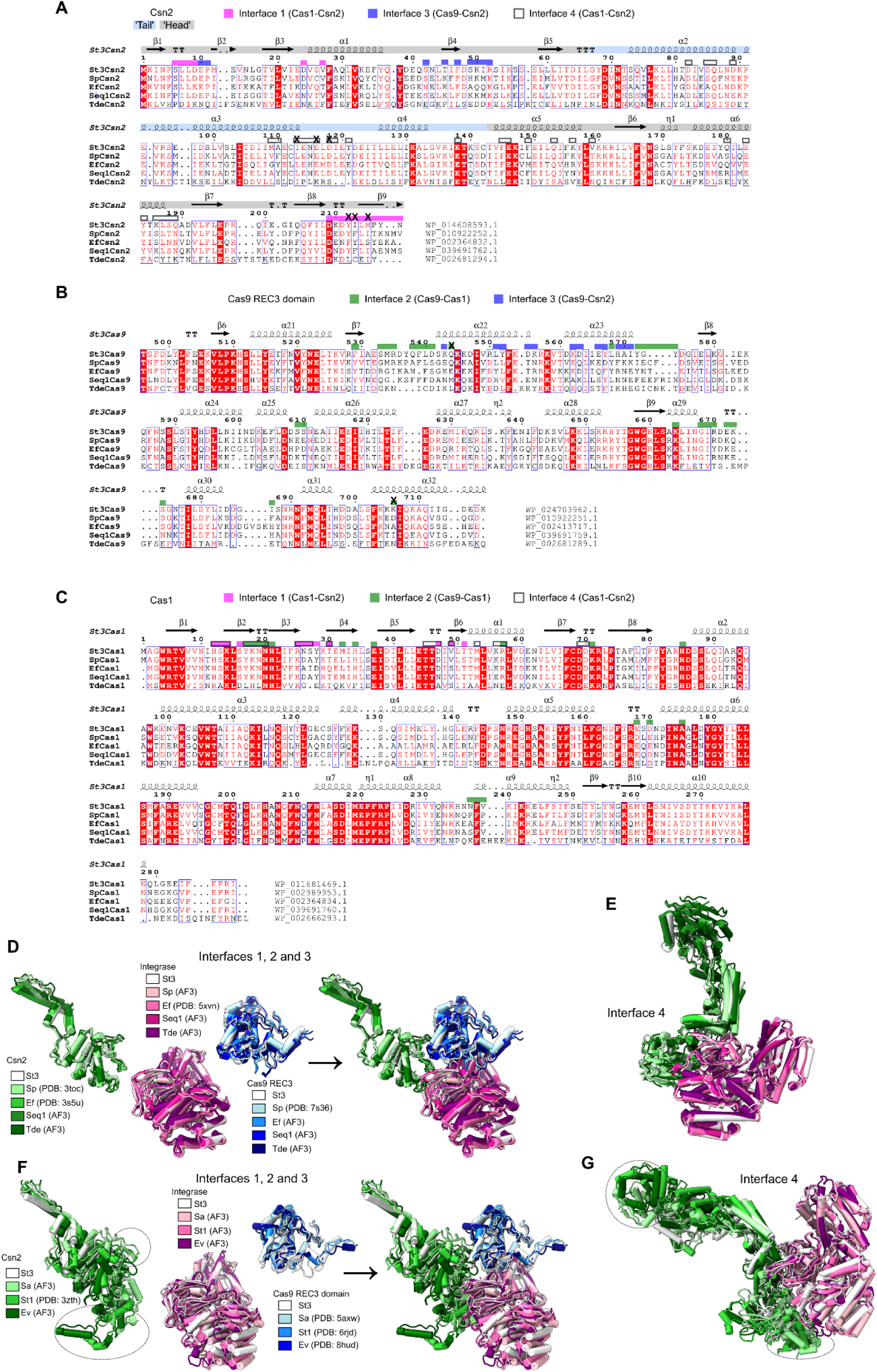
Comparison of type II-A Cas9, Cas1 and Csn2 proteins. (A)-(C) Sequence alignments of Csn2, Cas9 REC3 domain and Cas1 proteins from St3, *S. pyogenes* (Sp), *E. faecalis* (Ef), *Streptococcus equinus* (Seq1) and *Treponema denticola* (Tde). Secondary structure elements above alignments are marked based on PDB: 9hp9 chain L (St3Csn2), PDB: 9h2g (St3Cas9), and 9h1v chain B (St3Cas1), respectively. Residues of St3 proteins forming Interfaces 1-4 are marked by colored blocks. Residues that were mutated in this study are marked by ‘X’. St3Csn2 ‘head’ and ‘tail’ subdomains are indicated by light gray and light blue bands, respectively. (D-E) Structural comparison of Csn2, Cas1-Cas2 integrase and Cas9 proteins from St3 and other type II-A systems encoding St3Cas9-like Cas9 nucleases. (F-G) Structural comparison of Csn2, Cas1-Cas2 integrase and Cas9 proteins from St3 system and type II-A systems encoding Cas9 nucleases from the more divergent SaCas9 (*S. aureus*) and St1Cas9 (*S. thermophilus* DGCC 7710 CRISPR1) clade^40^. Overlays of Csn2, Cas1-Cas2 integrase and Cas9 proteins from other type II-A systems on the St3 supercomplex ((D) and (F)) and the St3 Interface 4-forming subcomplex ((E) and (G)) are shown. Experimental structures (where available) and AlphaFold 3^67^ models (‘AF3’) were used. For clarity, only the interface-forming subunits or domains are shown: Cas9 REC3 domain, a single subunit of Csn2 tetramer, and a single Cas1 dimer in (D); two Csn2 subunits and a single Cas1 dimer in (E). Components of the St3 supercomplex are white, proteins from other systems are colored in shades of pink (Cas1), blue (Cas9) and green (Csn2). The EfCsn2 (PDB: 3s5u) sequence is 99% identical to EfCsn2 encoded with the structurally characterized Ef integrase (PDB: 5xvn). Sequence of the *S. thermophilus* Csn2 with the available structure (‘StCsn2’, PDB: 3zth) is 97% identical to Csn2 from the *S. thermophilus* DGCC 7710 CRISPR1 system encoding St1Cas9 (PDB: 6rjc). The respective interaction surfaces in the *S. aureus*-like systems may be altered to accommodate additional Csn2 insertions in Sa/St/Ev Csn2 proteins not present in St3Csn2 (marked by dotted circles) and Cas9 REC domain reduction. EfCsn2 (PDB: 3s5u) shown in (D) has 99% sequence identity to EfCsn2 used for the alignment in (A).

**Figure S8.**
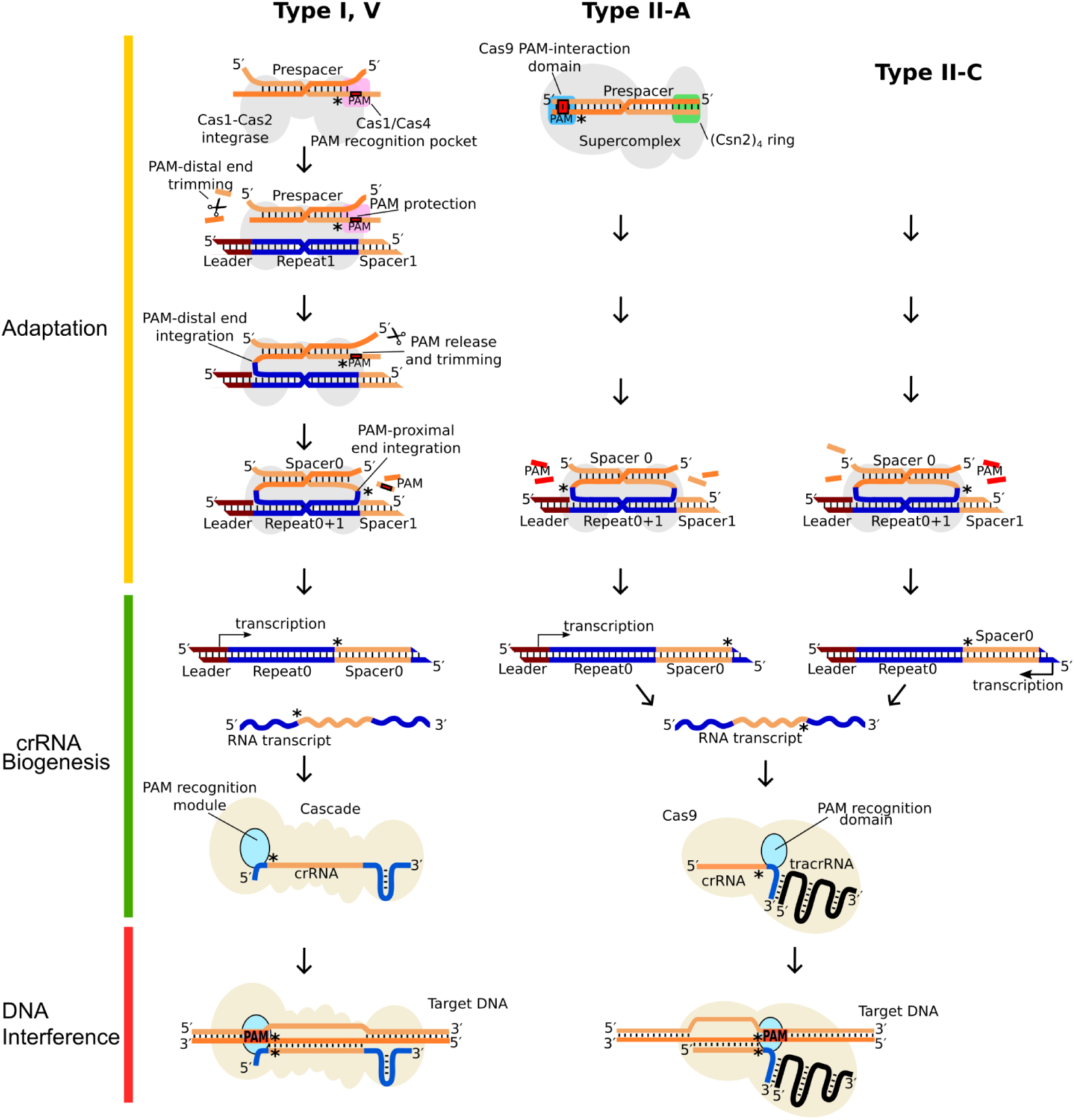
CRISPR immunity in type I, V and type II CRISPR-Cas systems. In the characterized type I and V systems (left), prespacer selection, processing and integration are carried out in the context of the Cas1-Cas2 integrase, which integrates the PAM-proximal prespacer end at the Repeat1-Spacer1 junction, ensuring that its transcript forms a functional crRNA for the Cascade (shown at the bottom) or the Cas12 effector complex (not shown). In the type II-A St3 systems (center) the prespacer is selected by the supercomplex, in which the PAM-distal end is occluded by Csn2. The PAM-proximal prespacer end in this case is inserted at the Leader-Repeat1 junction, ensuring formation of a functional Cas9:tracrRNA:crRNA complex (bottom). In the type II-C systems that lack Csn2 (right), the spacer is likely inserted in reverse orientation compared to type II-A systems. A functional Cas9 crRNA in this case is generated due to the reverse transcription direction of type II-C CRISPR region^43,44^. The asterisks (*) mark the PAM-proximal (pre)spacer and crRNA ends. Type II-B systems likely employ type I-like integration mechanism mediated by Cas1, Cas2 and Cas4, and type II-C-like transcription directionality or the CRISPR region to generate a valid crRNA.

**Figure S9.**
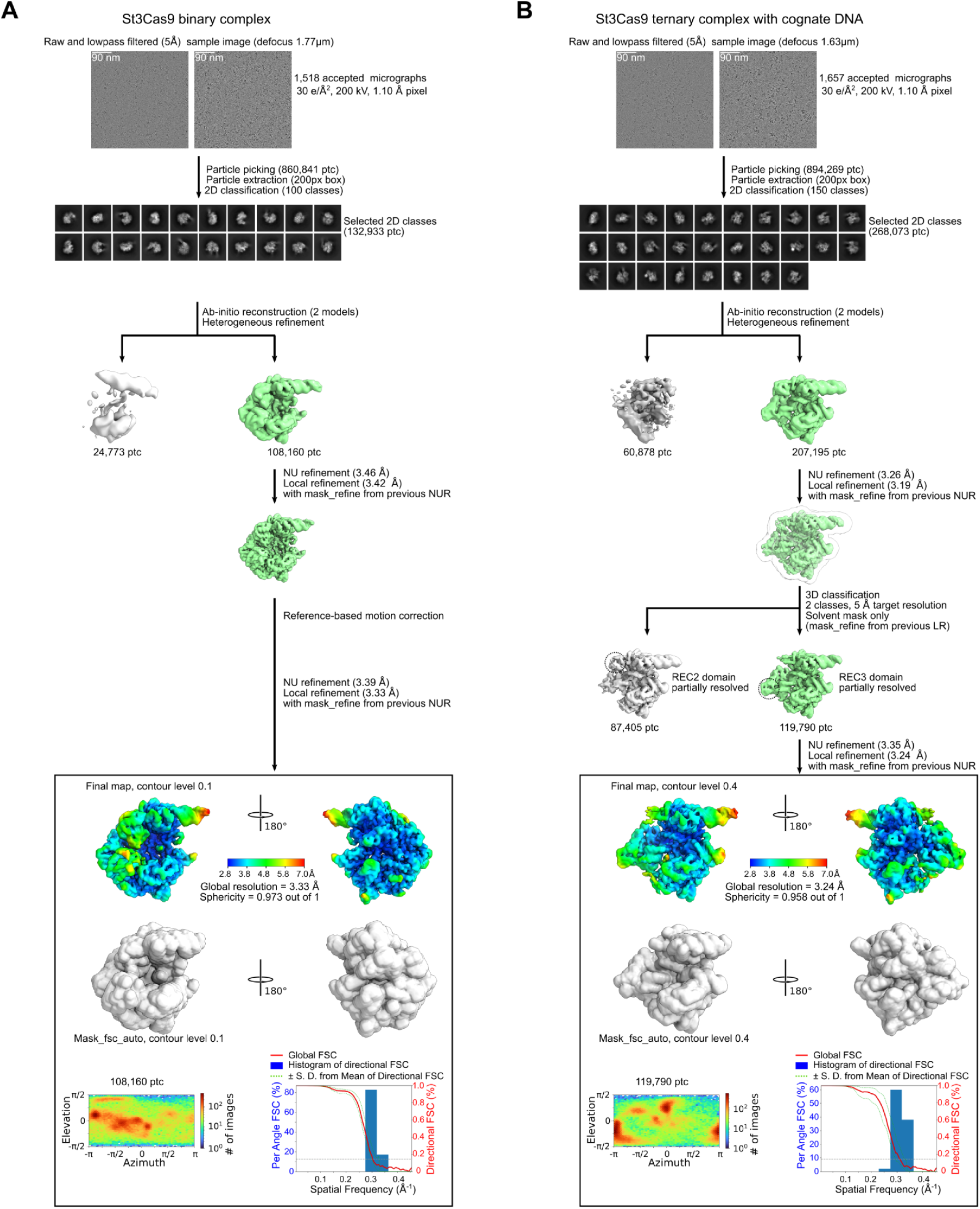
Processing of St3Cas9 cryo-EM data. (A) St3Cas9 binary complex. (B) St3Cas9 ternary complex with cognate DNA. ‘NU refinement’ (‘NUR’) denotes non-uniform refinement, ‘LR’ denotes local refinement.

**Figure S10.**
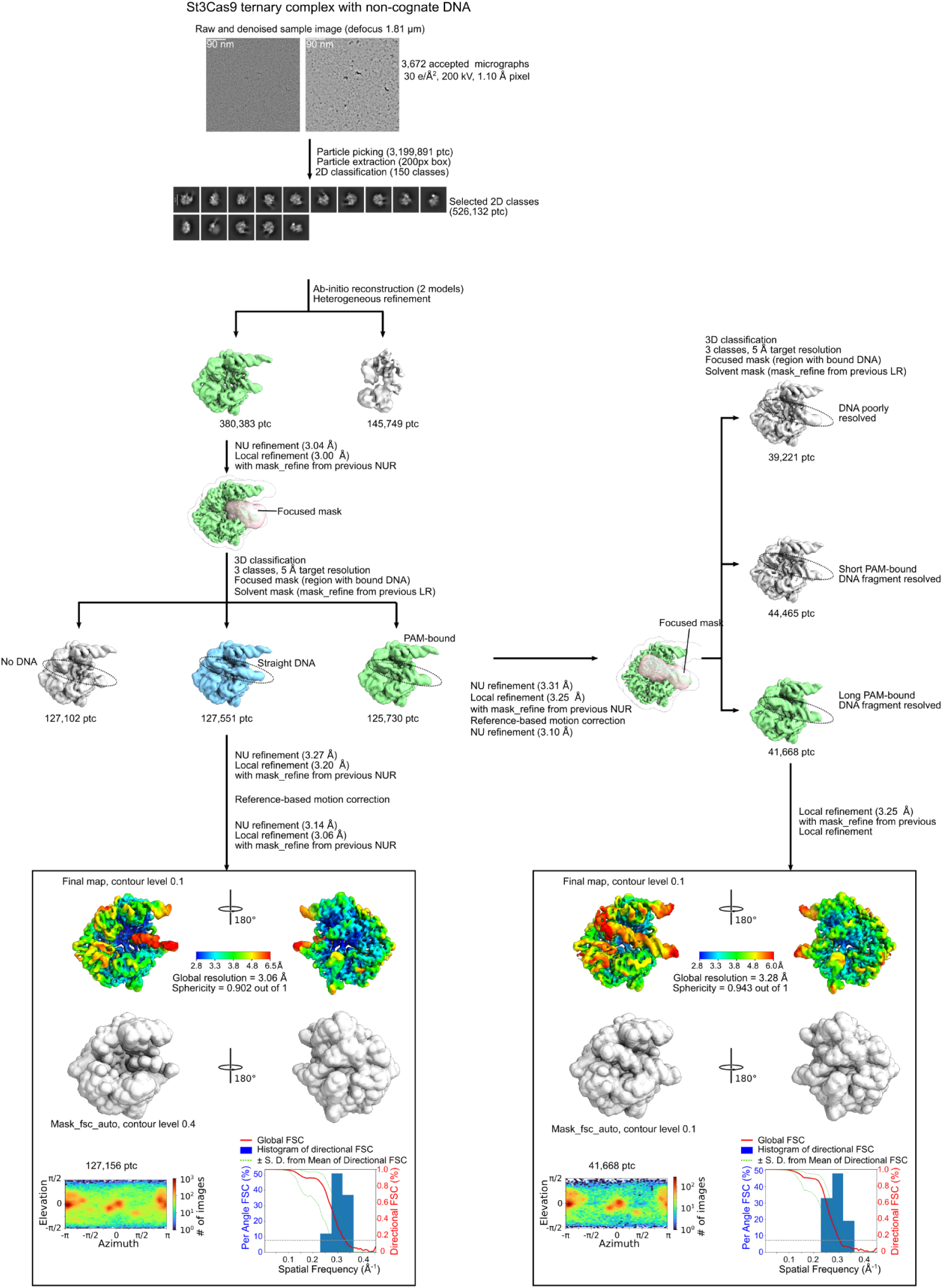
Processing of St3Cas9 ternary complex with non-cognate DNA cryo-EM data. ‘NU refinement’ (‘NUR’) denotes non-uniform refinement, ‘LR’ denotes local (focused) refinement.

**Figure S11.**
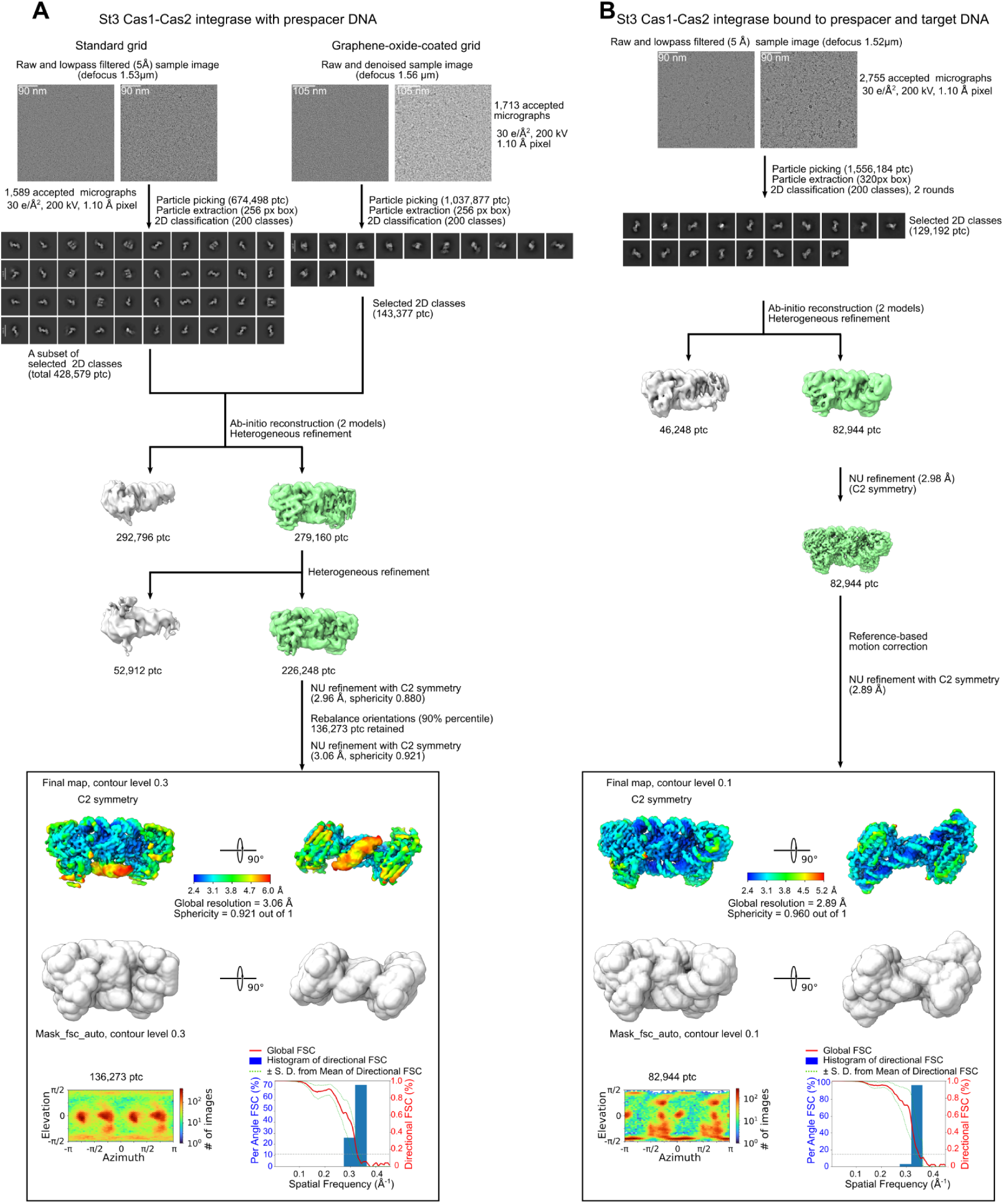
Processing of St3 Cas1-Cas2 integrase cryo-EM data. (A) St3 Cas1-Cas2 integrase bound to prespacer DNA. Particles extracted from micrographs obtained using standard and graphene oxide coated grids were combined after 2D classification. (B) St3 Cas1-Cas2 integrase bound to prespacer and target DNA. ‘NU refinement’ (‘NUR’) denotes non-uniform refinement.

**Figure S12.**
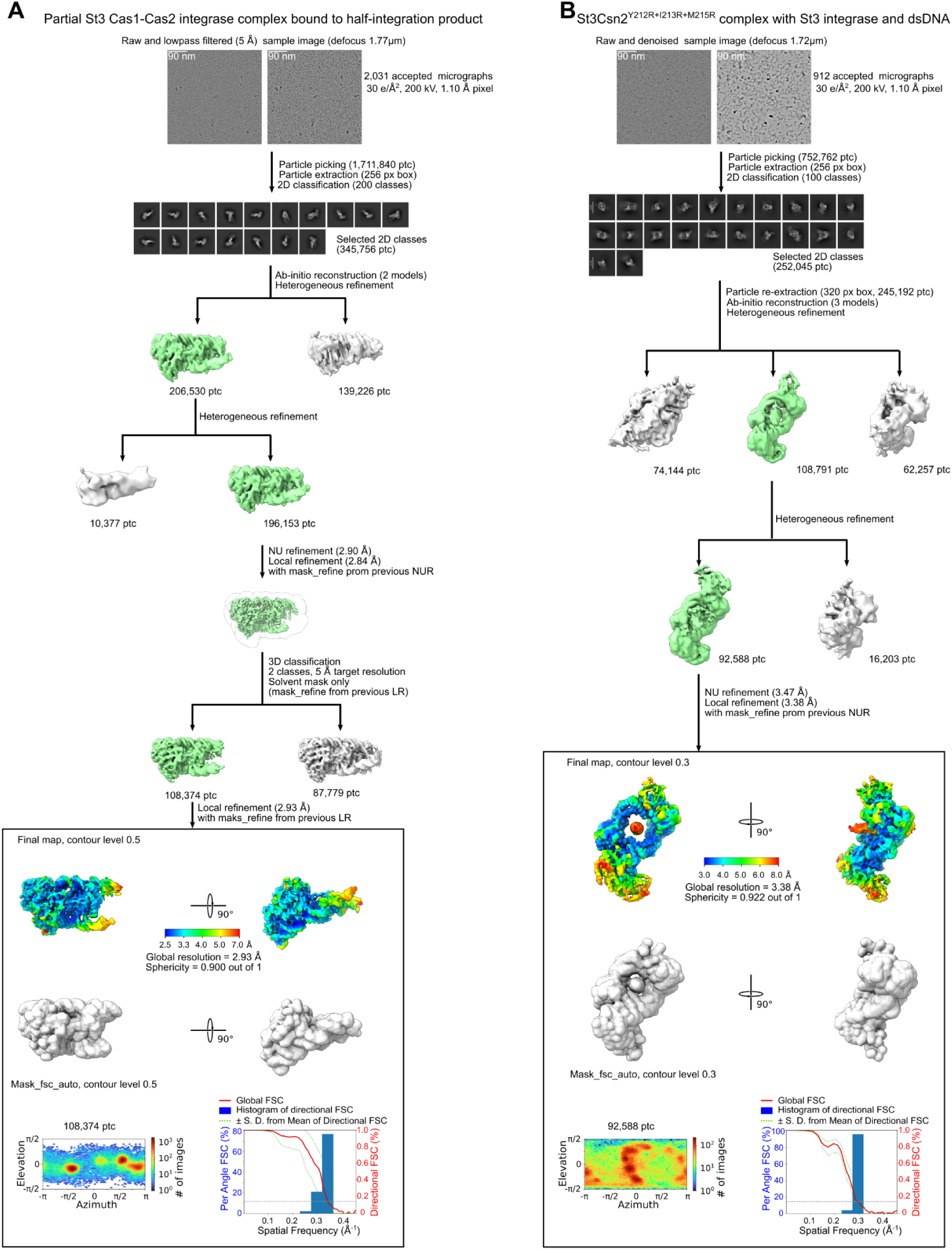
Cryo-EM data processing for the St3 spacer acquisition subcomplexes. (A) Partial St3 Cas1-Cas2 integrase bound to a half-integration product. (B) Csn2^Y212R+I213R+M215R^ complex with St3 integrase and dsDNA. ‘NU refinement’ and ‘NUR’ denote non-uniform refinement, ‘LR’ denotes local refinement.

**Figure S13.**
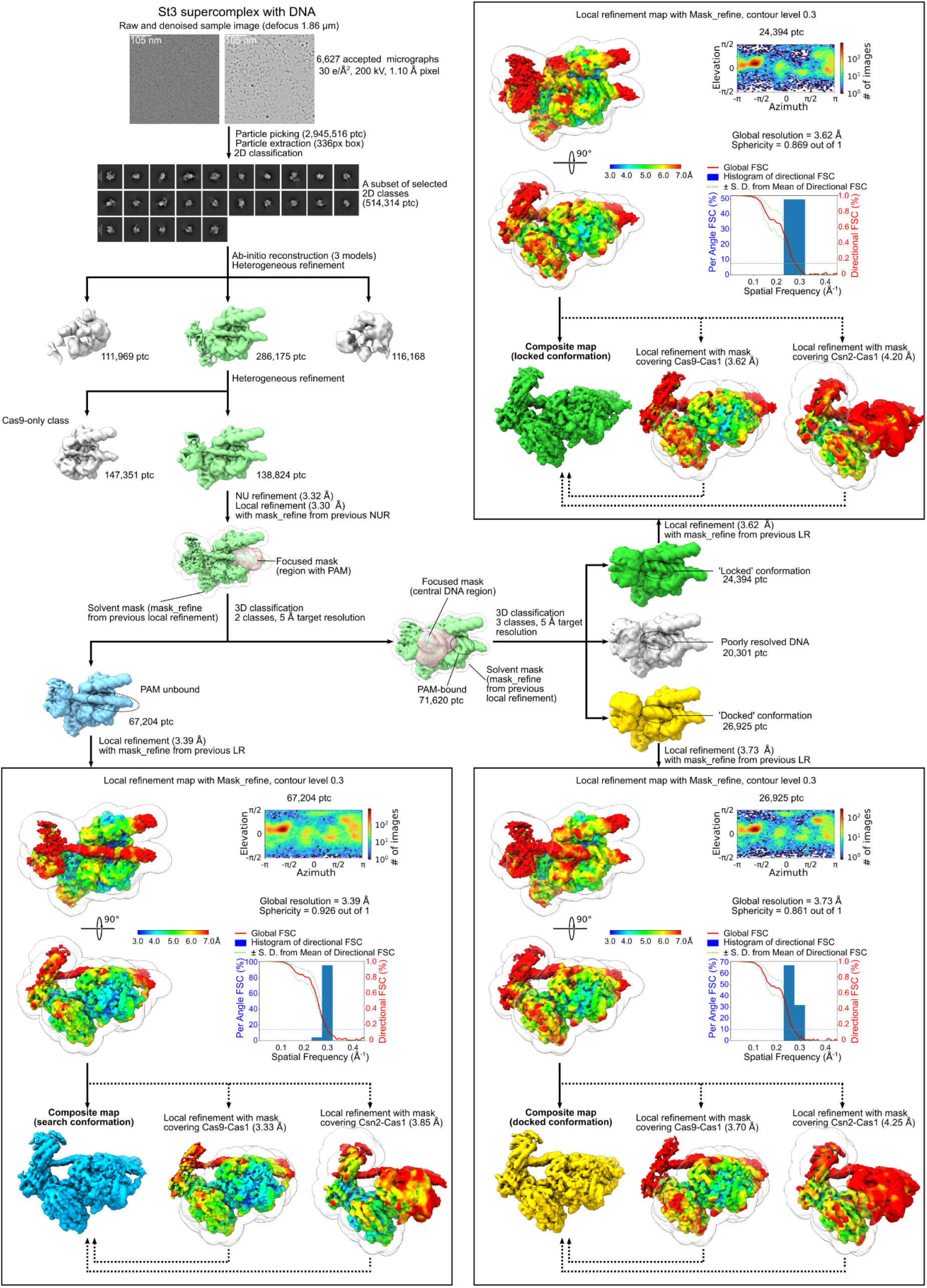
Processing of St3 supercomplex cryo-EM data. ‘NU refinement’ and ‘NUR’ denote non-uniform refinement, ‘LR’ denotes local refinement. 2D classes are also shown in Figure S3E.

**Movie S1. Cryo-EM maps and composition of the St3 supercomplex.** Unsharpened composite cryo-EM maps for the St3 supercomplex in three different conformations are shown: search (EMD-52324), docked (EMD-52320) and locked (EMD-52886) followed by the atomic model of the locked conformation (PDB: 9q85).

**Movie S2. Transition between the three different St3 supercomplex conformations.** The atomic models are morphed between the search (PDB: 9hp9), docked (PDB: 9hp8) and locked (PDB: 9q85) supercomplex conformations demonstrating the likely trajectory of captured DNA within the supercomplex during formation of the locked complex, and the associated rearrangement of the St3Cas9 REC2 domain.

**Table S1.**
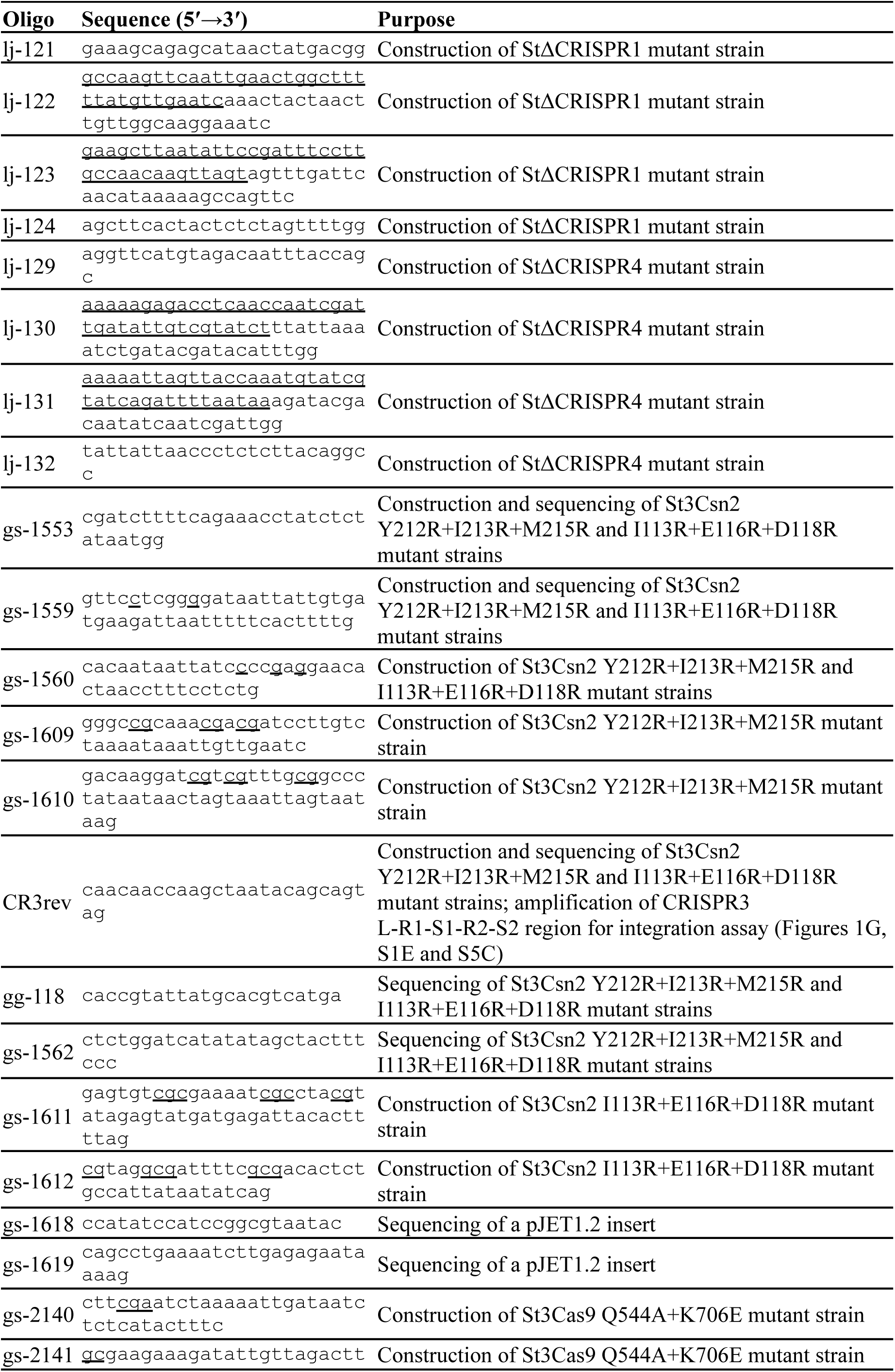
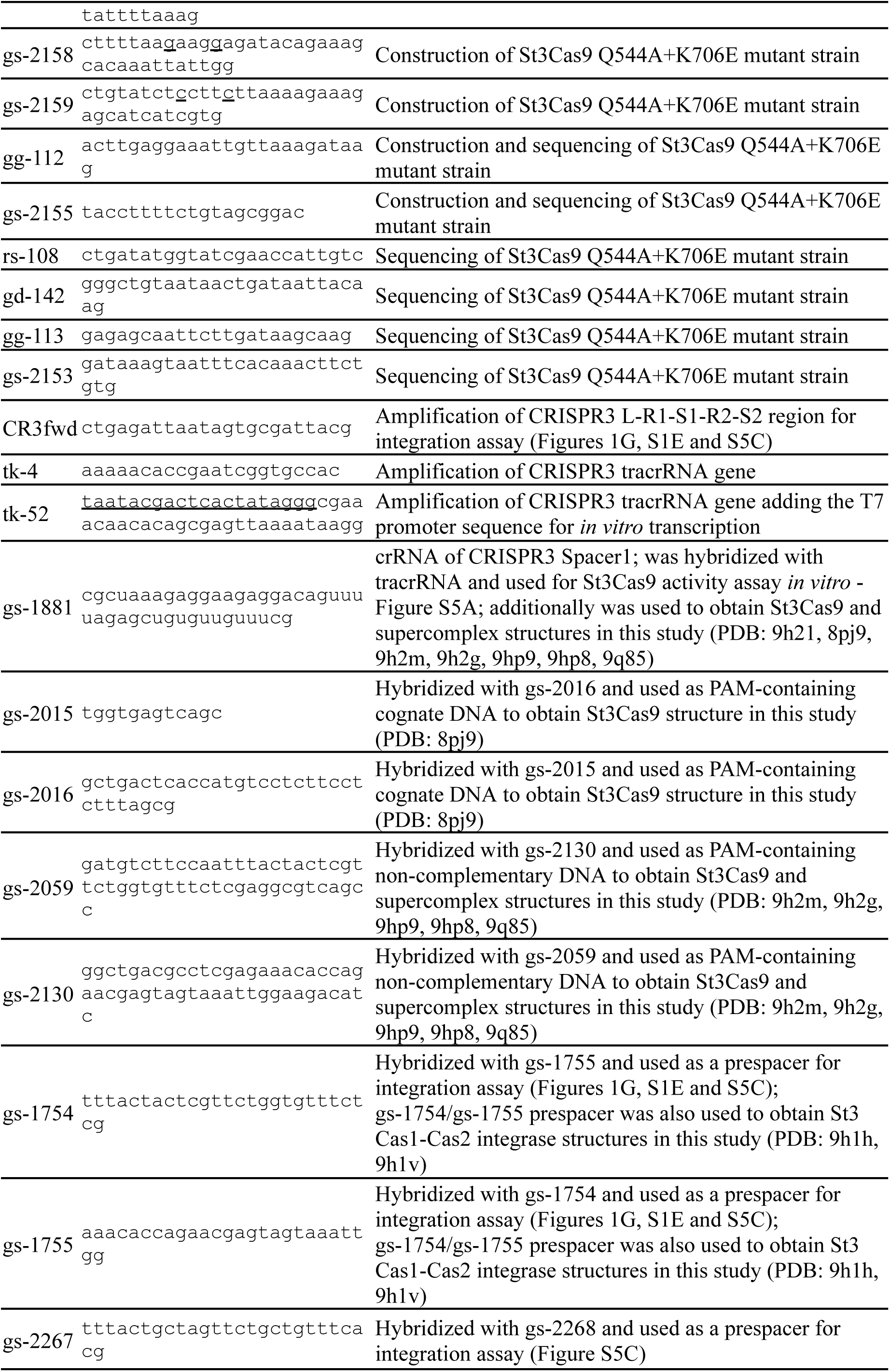
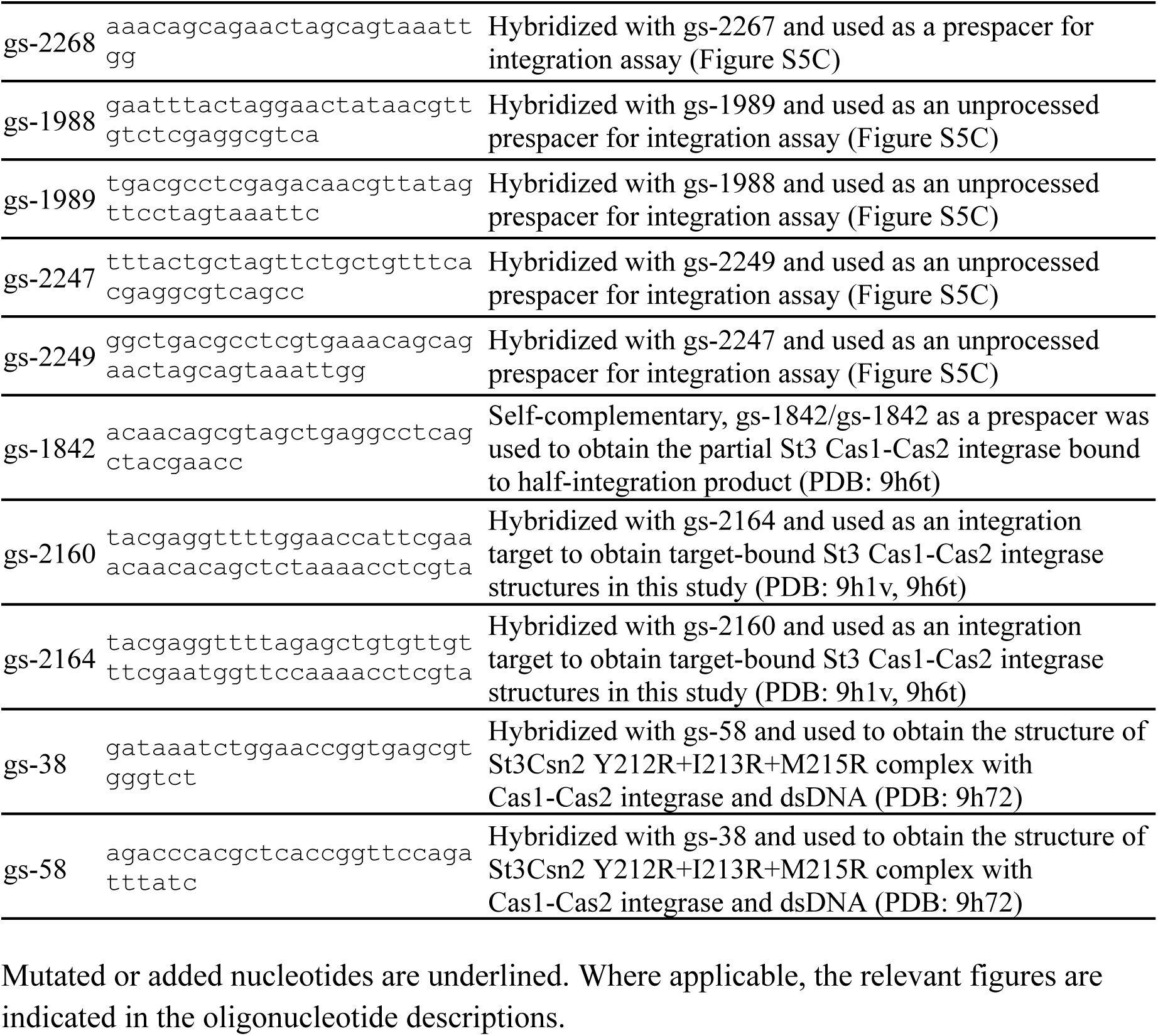
DNA oligonucleotides used in this study.

**Table S2.** Cryo-EM data collection, refinement and validation statistics.

|  | St3Cas9<br>binary<br>complex | St3Cas9 ternary complex |  |  | St3 Cas1-Cas2 integrase |  |  |  | St3 supercomplex * |  |  |
| --- | --- | --- | --- | --- | --- | --- | --- | --- | --- | --- | --- |
|  |  | Cognate DNA | Non-cognate DNA |  | Prespacer<br>DNA | Prespacer and<br>target DNA | Half<br>integration<br>product,<br>asymmetric<br>integrase<br>assembly | Csn2<sup>Y212R+I213R+</sup><br>M215R,<br>Cas1-Cas2<br>integrase and<br>dsDNA,<br>alternative<br>Cas1-Csn2<br>interaction<br>surface | Search<br>conformation | Bound PAM,<br>'docked'<br>conformation | Bound PAM,<br>'locked'<br>conformation |
|  |  |  | Unbound PAM | Bound PAM |  |  |  |  |  |  |  |
| PDB | 9h21 | 8pj9 | 9h2m | 9h2g | 9h1h | 9h1v | 9h6t | 9h72 | 9hp9 | 9hp8 | 9q85 |
| EMDB | EMD-51787 | EMD-17702 | EMD-51812 | EMD-51801 | EMD-51766 | EMD-51773 | EMD-51900 | EMD-51909 | EMD-52324<br>EMD-52321<br>EMD-52322<br>EMD-52323 | EMD-52320<br>EMD-52317<br>EMD-52318<br>EMD-52319 | EMD-52886<br>EMD-52883<br>EMD-52884<br>EMD-52885 |
| Data collection and processing |  |  |  |  |  |  |  |  |  |  |  |
| Hardware | Microscope: Glacios (TFS); voltage: 200 kV; detector: Falcon III camera (TFS); magnification: 92,000; automation software: EPU (TFS) |  |  |  |  |  |  |  |  |  |  |
| Electron<br>exposure<br>(e<sup>-</sup>/Å<sup>2</sup>) | 30 | 30 | 30 |  | 30 | 30 | 30 | 30 | 30 |  |  |
| Defocus range<br>(μm) | 0.5–2.5 | 0.5–2.5 | 0.5–2.5 |  | 0.5–2.5 | 0.5–2.5 | 0.5–2.5 | 0.5–2.5 | 0.5–2.5 |  |  |
| Pixel size (Å) | 1.1 | 1.1 | 1.1 |  | 1.1 | 1.1 | 1.1 | 1.1 | 1.1 |  |  |
| Micrographs<br>collected/used | 1,551/1,518 | 1,744/1,657 | 4,083/3,672 |  | 3,404/3,302 | 3,129/2,755 | 2,135/2,031 | 913/912 | 7,278/6,627 |  |  |
| Initial/final<br>particle<br>images (no.) | 860,841/<br>108,160 | 894,269/<br>119,790 | 3,199,891/<br>127,156 | 3,199,891/<br>41,668 | 1,712,375/<br>136,273 | 1,556,184/<br>82,944 | 1,711,840/<br>108,374 | 752,762/<br>92,588 | 2,945,516/<br>67,204 | 2,945,516/<br>26,925 | 2,945,516/<br>24,394 |
| Symmetry<br>imposed | C1 | C1 | C1 | C1 | C2 | C2 | C1 | C1 | C1 | C1 | C1 |
| Map<br>resolution (Å) | 3.33/4.0 | 3.24/4.1 | 3.06/3.9 | 3.28/4.2 | 3.06/3.4 | 2.89/3.4 | 2.93/4.1 | 3.38/6.2 | 3.39/6.4 * | 3.73/7.5 * | 3.62/7.6 * |
| (masked/unmasked); FSC threshold 0.143 |  |  |  |  |  |  |  |  |  |  |  |
| Map sharpening B factor ($\text{\AA}^2$) | 171.42 | 186.40 | 165.78 | 152.80 | 127.58 | 79.69 | 92.69 | 94.5 | 152.61 * | 186.64 * | 182.71 * |
| 3D FSC sphericity | 0.973 | 0.958 | 0.902 | 0.943 | 0.921 | 0.960 | 0.900 | 0.922 | 0.926 * | 0.861 * | 0.869 * |
| Refinement (Phenix) |  |  |  |  |  |  |  |  |  |  |  |
| Initial model used | AlphaFold 2 | AlphaFold 2 | 9h2g | AlphaFold 2 | AlphaFold 2 | AlphaFold 2 | 9h1v | 9h1v, AlphaFold 2 | 9h2m, 9h1v, AlphaFold 2 | 9h2g, 9h1v, AlphaFold 2 | 9hp8 |
| Model resolution ($\text{\AA}$)<br>FSC threshold 0.5 | 3.5 | 3.4 | 3.6 | 3.7 | 3.2 | 3.1 | 3.2 | 3.7 | 3.7 | 4.0 | 3.9 |
| Model composition |  |  |  |  |  |  |  |  |  |  |  |
| Non-hydrogen atoms | 12,065 | 11,163 | 13,211 | 13,185 | 11,628 | 13,773 | 7,841 | 11,094 | 23,569 | 23,090 | 23,107 |
| Protein residues | 1,336 | 1,051 | 1,337 | 1,366 | 1,360 | 1,372 | 768 | 1460 | 2720 | 2717 | 2702 |
| Nucleotides | 84 | 143 | 145 | 139 | 76 | 148 | 98 | 24 | 184 | 184 | 184 |
| Ligands (Ca<sup>2+</sup>) | 1 | 1 | 1 | 1 | 2 | 2 | 2 | 4 | 1 | 1 | 1 |
| B factors ($\text{\AA}^2$) min/max/mean | | | | | | | | | | | |
| Protein | 24.81/<br>203.95/<br>92.88 | 22.92/<br>167.67/<br>76.57 | 33.22/<br>206.56/<br>99.16 | 17.38/<br>203.62/<br>83.60 | 21.53/<br>215.29/<br>89.84 | 40.05/<br>174.33/<br>92.43 | 26.45/<br>160.92/<br>78.09 | 17.17/<br>236.51/<br>108.22 | 4.21/<br>312.34/<br>89.87 | 16.62/<br>275.26/<br>74.76 | 5.20/<br>339.00/<br>72.61 |
| Nucleotide | 25.56/<br>175.26/<br>76.85 | 3.40/<br>191.92/<br>84.06 | 2.38/<br>193.67/<br>90.34 | 3.21/<br>235.32/<br>95.36 | 0.04/<br>154.74/<br>48.57 | 5.92/<br>72.33/<br>34.39 | 1.32/<br>110.65/<br>33.22 | 82.99/<br>187.95/<br>130.73 | 16.85/<br>243.56/<br>110.97 | 11.44/<br>709.48/<br>138.49 | 12.02/<br>286.96/<br>86.59 |
| Ligands (Ca<sup>2+</sup>) | 52.12/<br>52.12/<br>52.12 | 25.89/<br>25.89/<br>25.89 | 48.05/<br>48.05/<br>48.05 | 41.13/<br>41.13/<br>41.13 | 32.57/<br>34.95/<br>33.76 | 54.25/<br>57.64/<br>55.94 | 49.39/<br>51.90/<br>50.64 | 86.79/<br>137.09/<br>105.19 | 24.43/<br>24.43/<br>24.43 | 61.20/<br>61.20/<br>61.20 | 51.72/<br>51.72/<br>51.72 |

| RMSDs |  |  |  |  |  |  |  |  |  |  |  |
| --- | --- | --- | --- | --- | --- | --- | --- | --- | --- | --- | --- |
| Bond lengths (Å) | 0.003 | 0.003 | 0.003 | 0.002 | 0.003 | 0.004 | 0.004 | 0.005 | 0.002 | 0.004 | 0.004 |
| Bond angles (°) | 0.528 | 0.664 | 0.701 | 0.477 | 0.611 | 0.605 | 0.690 | 0.757 | 0.414 | 0.707 | 0.690 |
| Validation |  |  |  |  |  |  |  |  |  |  |  |
| MolProbity score | 1.38 | 1.64 | 1.42 | 1.47 | 1.60 | 1.71 | 2.03 | 2.20 | 1.36 | 1.90 | 1.94 |
| Clashscore | 6 | 13 | 8 | 9 | 8 | 6 | 9 | 11 | 6 | 12 | 11 |
| Poor rotamers (%) | 1.2 | 1.2 | 0.9 | 1.0 | 1.9 | 1.7 | 2.1 | 1.8 | 1.1 | 2.3 | 1.5 |
| Ramachandran plot |  |  |  |  |  |  |  |  |  |  |  |
| Favored (%) | 98.05 | 99.14 | 98.35 | 98.53 | 97.26 | 98.24 | 97.10 | 96.20 | 98.33 | 97.51 | 96.93 |
| Allowed (%) | 1.95 | 0.86 | 1.65 | 1.47 | 2.74 | 1.76 | 2.90 | 3.80 | 1.67 | 2.49 | 3.07 |
| Disallowed (%) | 0 | 0 | 0 | 0 | 0 | 0 | 0 | 0 | 0 | 0 | 0 |
| CCvolume/CC mask | 0.79/0.80 | 0.76/0.78 | 0.64/0.65 | 0.71/0.73 | 0.74/0.75 | 0.80/0.80 | 0.66/0.66 | 0.76/0.77 | 0.71/0.73 | 0.71/0.72 | 0.72/0.73 |
| Cβ outliers (%) | 0 | 0 | 0 | 0 | 0 | 0 | 0 | 0 | 0 | 0 | 0 |
| CaBLAM outliers (%) | 1.89 | 1.94 | 0.98 | 0.96 | 0.97 | 0.89 | 1.20 | 1.75 | 1.38 | 1.27 | 1.47 |

\* - For the supercomplexes, four EMDB codes are shown: the composite map, the consensus map, focused map 1 (Cas1-Cas9 mask), and focused map 2 (Cas1-Csn2 mask); map resolution and sphericity score are reported for the consensus map; map sharpening B factor is reported for the composite map.

## REFERENCES

1. Barrangou, R. et al. CRISPR provides acquired resistance against viruses in prokaryotes. Science 315, 1709–1712 (2007).

2. Nuñez, J. K., Harrington, L. B., Kranzusch, P. J., Engelman, A. N. & Doudna, J. A. Foreign DNA capture during CRISPR-Cas adaptive immunity. Nature 527, 535–538 (2015).

3. Yosef, I., Goren, M. G. & Qimron, U. Proteins and DNA elements essential for the CRISPR adaptation process in Escherichia coli. Nucleic Acids Res 40, 5569–5576 (2012).

4. Wei, Y., Chesne, M. T., Terns, R. M. & Terns, M. P. Sequences spanning the leader-repeat junction mediate CRISPR adaptation to phage in Streptococcus thermophilus. Nucleic Acids Research 43, 1749–1758 (2015).

5. Xiao, Y., Ng, S., Hyun Nam, K. & Ke, A. How type II CRISPR-Cas establish immunity through Cas1-Cas2-mediated spacer integration. Nature 550, 137–141 (2017).

6. Wright, A. V. & Doudna, J. A. Protecting genome integrity during CRISPR immune adaptation. Nat Struct Mol Biol 23, 876–883 (2016).

7. Jackson, S. A. et al. CRISPR-Cas: Adapting to change. Science 356, eaal5056 (2017).

8. Mojica, F. J., Diez-Villasenor, C., Garcia-Martinez, J. & Almendros, C. Short motif sequences determine the targets of the prokaryotic CRISPR defence system. Microbiology 155, (2009).

9. Anders, C., Niewoehner, O., Duerst, A. & Jinek, M. Structural basis of PAM-dependent target DNA recognition by the Cas9 endonuclease. Nature 513, 569–573 (2014).

10. Marraffini, L. A. & Sontheimer, E. J. Self versus non-self discrimination during CRISPR RNA-directed immunity. Nature 463, 568–571 (2010).

11. Sasnauskas, G. & Siksnys, V. CRISPR adaptation from a structural perspective. Curr Opin Struct Biol 65, 17–25 (2020).

12. Lee, H. & Sashital, D. G. Creating memories: molecular mechanisms of CRISPR adaptation. Trends in Biochemical Sciences 47, 464–476 (2022).

13. Hu, C. et al. Mechanism for Cas4-assisted directional spacer acquisition in CRISPR-Cas. Nature 598, 515–520 (2021).

14. Hudaiberdiev, S. Phylogenomics of Cas4 family nucleases. BMC Evol. Biol. 17, (2017).

15. Dhingra, Y., Suresh, S. K., Juneja, P. & Sashital, D. G. PAM binding ensures orientational integration during Cas4-Cas1-Cas2-mediated CRISPR adaptation. Mol Cell 82, 4353–4367.e6 (2022).

16. Lee, H., Zhou, Y., Taylor, D. W. & Sashital, D. G. Cas4-Dependent Prespacer Processing Ensures High-Fidelity Programming of CRISPR Arrays. Mol Cell 70, 48–59.e5 (2018).

17. Lee, H., Dhingra, Y. & Sashital, D. G. The Cas4-Cas1-Cas2 complex mediates precise prespacer processing during CRISPR adaptation. eLife 8, (2019).

18. Kieper, S. N. et al. Cas4 Facilitates PAM-Compatible Spacer Selection during CRISPR Adaptation. Cell Rep 22, 3377–3384 (2018).

19. Shiimori, M., Garrett, S. C., Graveley, B. R. & Terns, M. P. Cas4 Nucleases Define the PAM, Length, and Orientation of DNA Fragments Integrated at CRISPR Loci. Molecular Cell 70, 814–824.e6 (2018).

20. Wang, J. et al. Structural and Mechanistic Basis of PAM-Dependent Spacer Acquisition in CRISPR-Cas Systems. Cell 163, 840–853 (2015).

21. Wang, J. Y. et al. Genome expansion by a CRISPR trimmer-integrase. Nature 618, 855–861 (2023).

22. Kim, S. et al. Selective loading and processing of prespacers for precise CRISPR adaptation. Nature 579, 141–145 (2020).

23. Drabavicius, G. et al. DnaQ exonuclease-like domain of Cas2 promotes spacer integration in a type I-E CRISPR-Cas system. EMBO Rep 19, e45543 (2018).

24. Ramachandran, A., Summerville, L., Learn, B. A., DeBell, L. & Bailey, S. Processing and integration of functionally oriented prespacers in the Escherichia coli CRISPR system depends on bacterial host exonucleases. J Biol Chem 295, 3403–3414 (2020).

25. Yoganand, K. N., Muralidharan, M., Nimkar, S. & Anand, B. Fidelity of prespacer capture and processing is governed by the PAM-mediated interactions of Cas1-2 adaptation complex in CRISPR-Cas type I-E system. Journal of Biological Chemistry 294, 20039–20053 (2019).

26. Heler, R. et al. Cas9 specifies functional viral targets during CRISPR–Cas adaptation. Nature 519, 199–202 (2015).

27. Wei, Y., Terns, R. M. & Terns, M. P. Cas9 function and host genome sampling in Type II-A CRISPR–Cas adaptation. Genes Dev. 29, 356–361 (2015).

28. Makarova, K. S. et al. Evolutionary classification of CRISPR-Cas systems: a burst of class 2 and derived variants. Nat Rev Microbiol 18, 67–83 (2020).

29. Ka, D., Jang, D. M., Han, B. W. & Bae, E. Molecular organization of the type II-A CRISPR adaptation module and its interaction with Cas9 via Csn2. Nucleic Acids Research 46, 9805–9815 (2018).

30. Wilkinson, M. et al. Structure of the DNA-Bound Spacer Capture Complex of a Type II CRISPR-Cas System. Molecular cell 75, 90–101.e5 (2019).

31. Jakhanwal, S. et al. A CRISPR-Cas9-integrase complex generates precise DNA fragments for genome integration. Nucleic Acids Res 49, 3546–3556 (2021).

32. Horvath, P. et al. Diversity, activity, and evolution of CRISPR loci in Streptococcus thermophilus. J Bacteriol 190, 1401–1412 (2008).

33. Glemzaite, M. et al. Targeted gene editing by transfection of in vitro reconstituted Streptococcus thermophilus Cas9 nuclease complex. RNA Biol 12, 1–4 (2015).

34. Cofsky, J. C., Soczek, K. M., Knott, G. J., Nogales, E. & Doudna, J. A. CRISPR–Cas9 bends and twists DNA to read its sequence. Nat Struct Mol Biol 29, 395–402 (2022).

35. Hibshman, G. N. et al. Unraveling the mechanisms of PAMless DNA interrogation by SpRY-Cas9. Nat Commun 15, 3663 (2024).

36. Arslan, Z. et al. Double-strand DNA end-binding and sliding of the toroidal CRISPR-associated protein Csn2. Nucleic Acids Research 41, 6347 (2013).

37. Moldovan, G.-L., Pfander, B. & Jentsch, S. PCNA, the Maestro of the Replication Fork. Cell 129, 665–679 (2007).

38. Hofmann, R., Herman, C., Mo, C. Y., Mathai, J. & Marraffini, L. A. Deep mutational scanning identifies variants of Cas1 and Cas2 that increase spacer acquisition in type II-A CRISPR-Cas systems. Preprint at 10.1101/2024.10.10.617623 (2024).

39. Chylinski, K., Makarova, K. S., Charpentier, E. & Koonin, E. V. Classification and evolution of type II CRISPR-Cas systems. Nucleic Acids Research 42, 6091–6105 (2014).

40. Gasiunas, G. et al. A catalogue of biochemically diverse CRISPR-Cas9 orthologs. Nat Commun 11, 5512 (2020).

41. Bestas, B. et al. A Type II-B Cas9 nuclease with minimized off-targets and reduced chromosomal translocations in vivo. Nat Commun 14, 5474 (2023).

42. Chen, F. et al. Targeted activation of diverse CRISPR-Cas systems for mammalian genome editing via proximal CRISPR targeting. Nat Commun 8, 14958 (2017).

43. Zhang, Y. et al. Processing-independent CRISPR RNAs limit natural transformation in Neisseria meningitidis. Mol Cell 50, 488–503 (2013).

44. Dugar, G. et al. High-Resolution Transcriptome Maps Reveal Strain-Specific Regulatory Features of Multiple Campylobacter jejuni Isolates. PLoS Genet 9, e1003495 (2013).

45. Workman, R. E. et al. A natural single-guide RNA repurposes Cas9 to autoregulate CRISPR-Cas expression. Cell 184, 675–688.e19 (2021).

46. Strecker, J. et al. RNA-guided DNA insertion with CRISPR-associated transposases. Science 365, 48–53 (2019).

47. Halpin-Healy, T. S., Klompe, S. E., Sternberg, S. H. & Fernández, I. S. Structural basis of DNA targeting by a transposon-encoded CRISPR–Cas system. Nature 577, 271–274 (2020).

48. Querques, I., Schmitz, M., Oberli, S., Chanez, C. & Jinek, M. Target site selection and remodelling by type V CRISPR-transposon systems. Nature 599, 497–502 (2021).

49. Jinek, M. et al. A programmable dual-RNA-guided DNA endonuclease in adaptive bacterial immunity. Science 337, 816–821 (2012).

50. Gasiunas, G., Barrangou, R., Horvath, P. & Siksnys, V. Cas9-crRNA ribonucleoprotein complex mediates specific DNA cleavage for adaptive immunity in bacteria. Proceedings of the National Academy of Sciences of the United States of America 109, (2012).

51. Shipman, S. L., Nivala, J., Macklis, J. D. & Church, G. M. Molecular recordings by directed CRISPR spacer acquisition. Science 353, aaf1175 (2016).

52. Fontaine, L. et al. Development of a Versatile Procedure Based on Natural Transformation for Marker-Free Targeted Genetic Modification in Streptococcus thermophilus. Applied and Environmental Microbiology 76, 7870–7877 (2010).

53. Labrie, S. J. et al. A mutation in the methionine aminopeptidase gene provides phage resistance in Streptococcus thermophilus. Sci Rep 9, 13816 (2019).

54. Hynes, A. P. et al. Detecting natural adaptation of the Streptococcus thermophilus CRISPR-Cas systems in research and classroom settings. Nat Protoc 12, 547–565 (2017).

55. Sapranauskas, R. et al. The Streptococcus thermophilus CRISPR/Cas system provides immunity in Escherichia coli. Nucleic Acids Research 39, 9275–9282 (2011).

56. Bokori-Brown, M. et al. Cryo-EM structure of lysenin pore elucidates membrane insertion by an aerolysin family protein. Nat Commun 7, 11293 (2016).

57. Punjani, A., Rubinstein, J. L., Fleet, D. J. & Brubaker, M. A. cryoSPARC: algorithms for rapid unsupervised cryo-EM structure determination. Nat Methods 14, 290–296 (2017).

58. Punjani, A., Zhang, H. & Fleet, D. J. Non-uniform refinement: adaptive regularization improves single-particle cryo-EM reconstruction. Nat Methods 17, 1214–1221 (2020).

59. Tan, Y. Z. et al. Addressing preferred specimen orientation in single-particle cryo-EM through tilting. Nat Methods 14, 793–796 (2017).

60. Liebschner, D. et al. Macromolecular structure determination using X-rays, neutrons and electrons: recent developments in Phenix. Acta Cryst D 75, 861–877 (2019).

61. Jumper, J. et al. Highly accurate protein structure prediction with AlphaFold. Nature 596, 583–589 (2021).

62. Pettersen, E. F. et al. UCSF ChimeraX: Structure visualization for researchers, educators, and developers. Protein Sci 30, 70–82 (2021).

63. Emsley, P., Lohkamp, B., Scott, W. G. & Cowtan, K. Features and development of Coot. Acta Crystallogr D Biol Crystallogr 66, 486–501 (2010).

64. Olechnovič, K. & Venclovas, Č. VoroContacts: a tool for the analysis of interatomic contacts in macromolecular structures. Bioinformatics 37, 4873–4875 (2021).

65. Madeira, F. et al. The EMBL-EBI Job Dispatcher sequence analysis tools framework in 2024. Nucleic Acids Res 52, W521–W525 (2024).

66. Robert, X. & Gouet, P. Deciphering key features in protein structures with the new ENDscript server. Nucleic Acids Research 42, W320–W324 (2014).

67. Abramson, J. et al. Accurate structure prediction of biomolecular interactions with AlphaFold 3. Nature 1–3 (2024) doi:10.1038/s41586-024-07487-w.

